# Uncovering the Biological Function of the Peptidoglycan Hydrolase PcsB in Streptococcus pneumoniae

**DOI:** 10.1101/2025.05.23.655736

**Authors:** Nicholas S Briggs, Kevin E Bruce, Stefano Camaione, Souvik Naskar, Adrian Lloyd, Malcolm E Winkler, David I Roper

**Author notes:** To whom correspondence should be addressed. Nicholas S Briggs, School of Life Sciences, University of Warwick, CV4 7AL, & David I Roper, School of Life Sciences, University of Warwick, CV4 7AL. **Significance Statement** Bacterial cell division requires coordinated assembly of specific proteins acting on the cell envelope to facilitate cell separation. In *Streptococcus pneumoniae*, the cell division complex FtsEX is recruited early in divisome assembly and controls the extracellular hydrolase function of PcsB, which cleaves linkages in the peptidoglycan layer. Here we demonstrate the connection between ATP hydrolysis in cytosolic FtsE, mechanotransduction through FtsX in the membrane, activation of PcsB in the intracellular space and molecular details of the chemical modification of the peptidoglycan layer as a result of these essential pneumococcal proteins. In addition, we present a model whereby the FtsEX-PcsB complex is assembled early in order to produce a hydrolase-modified form of peptidoglycan during septal splitting, marking the site of later division events, thus providing a rationale for its biological and temporal function.

## Abstract

In the human commensal Gram-positive bacterial pathogen Streptococcus pneumoniae, the essential cell-division-associated peptidoglycan (PG) hydrolase PcsB interacts directly with the membrane-bound FtsEX complex (1-3). PcsB contains a cysteine, histidine-dependent amidohydrolase/peptidase (CHAP) domain responsible for PG hydrolysis, and a coiled-coil domain required for interaction with FtsEX (1,4). ATP hydrolysis of FtsE in the cytoplasm drives conformational changes in FtsX in the cytoplasmic membrane, which ultimately regulates the PG hydrolase on the outside of the cell (5). In this work we show that the CHAP domain of PcsB predominately functions as an D-iso-Glutaminyl-Lysyl D,L-endopeptidase, with substrate specificity for Lys-containing, amidated PG. The catalytic activity of PcsB is regulated by conformational changes of the coiled-coil region and by a short helical region immediately adjacent to the CHAP domain, to guard against PcsB hydrolytic activation outside of its cell division specific functional requirement. This work supports a model for the overall biological activity of the FtsEX-PcsB complex, allowing stem peptide cleavage that splits the septal disk and marks a region of the peptidoglycan sacculus for subsequent cell division remodelling.

## Introduction

Cell division represents one of the most fundamental and complex processes undertaken by a bacterial cell. Central to this process is cleavage of the bacterial cell wall structural polymer peptidoglycan to enable separation of individual cells (1,2). During cell division, selected conserved proteins dynamically facilitate alterations to membrane and cell wall morphology, enabling coordination between events located on both the inside and outside of the cytoplasmic membrane (2). Previous studies of bacterial cell morphology in Escherichia coli (3,4) and Bacillus subtilis (5–7) in particular, have identified key proteins and complexes required for division using detailed genetic, biochemical, and advanced microscopy techniques. Homologues of many of the key cell division proteins are found in almost all bacterial species, that importantly underlines their essentiality and common genetic origin (2,3,8,9).

In the early stages of bacterial cell division, a complex between the FtsE and FtsX (FtsEX) proteins is formed (10–13) and is recruited via interaction with the filament-forming proteins FtsA and FtsZ to form components of the cytoplasmic Z-ring (14,15). The Z-ring provides the foundation for the later assembly of a complex termed the bacterial divisome (2,3,16) which accomplishes the process of cell division. FtsX is an integral membrane protein which forms a dimer and partners with a dimer of FtsE, the cytoplasmic nucleotide-binding domain of the FtsEX complex (10–13,17,18). The FtsEX complex is an important member of the early divisome apparatus mediating important protein-protein interactions and also has a direct role in controlling PG degradation activity via extracellular hydrolytic proteins (11–13,17). Whilst FtsEX is a structural homologue of the type VII family of ATP dependent ABC transporters, it has no transport role but instead uses ATP hydrolysis in FtsE to transduce a signal through FtsX to extracellular partner proteins to bring about controlled PG degradation (11–13,17,18).

In rod-shaped Gram-negative bacteria including *E. coli*, FtsEX is not essential but is required for viability in low-osmolarity media (17). By contrast in the ovoid-shaped Gram-positive strain *Streptococcus pneumoniae,* FtsEX and PcsB are essential (*Spn* or pneumococcus) (1,3). The FtsEX protein complex is further delineated in Gram-positive and Gram-negative bacteria by the relationship to their cognate PG hydrolase enzymes which they activate and control. In typical Gram-negative bacterial species such as *E. coli*, the FtsEX protein complex binds to a structural, regulatory coiled-coil protein EnvC, which itself has no enzymatic activity, but binds to and regulates the activity of one or a series of zinc-dependent PG amidase enzymes (AmiA, AmiB) in order to degrade the PG (19,20). In Gram-positive bacteria such as *B. subtilis* and *S. pneumoniae,* the structural regulatory domain and enzymatic PG hydrolase are combined in a single protein such as CwlO (18,21) or PcsB (10,22) respectively, which interacts directly with the FtsEX complex. Why Gram-negative species like *E. coli* and *Pseudomonas aeruginosa* have alternative PG hydrolytic enzymes (AmiA/AmiB) whose association with FtsEX requires an adaptor protein (EnvC) and what determines their selection, remains an unexplored area in bacterial cell wall biology but may be linked to the greater compartmentalised structure of the Gram-negative cell envelope and its regulation and maintenance.

It has been shown previously that the full-length *S. pneumoniae* PcsB protein displays no hydrolytic activity *in vitro* until the N-terminal coiled-coil region of the protein is removed (22), suggesting a critical conformational regulatory function is required for PcsB PG hydrolytic activation. The remaining C-terminal section of PcsB is a cysteine, histidine-dependent amidohydrolase/peptidase (CHAP) domain that shares overall homology to other PG hydrolases (13,18,21), and thus was assumed to play a similar role to that of the *E. coli* amidases during cell division which target the MurNAc-L-Ala bond to produce fully denuded glycan strands (20,23). However, whilst PcsB has no metal dependency and has closer homology to other Gram-positive PG hydrolases like the *B. subtilis* D,L-endopeptidase CwlO (18), little is known about PcsB’s *in vivo* molecular target and cleavage product and how this relates to the essential role it plays during cell division.

The X-ray crystal structure of the full-length PcsB protein revealed a dimeric head-to-head arrangement of the CHAP domains that prevented PG degradation activity (22), but biochemical studies revealed the activity of the CHAP domain in isolation. Subsequent studies on the structures of *E. coli* EnvC-AmiA/B (20) as well as the complexes of *P. aeruginosa* FtsX-EnvC, *P. aeruginosa* FtsEX-EnvC-AmiB (24), *Vibrio cholerae* FtsEX-EnvC (12), *Mycobacterium tuberculosis* FtsEX-RipC (13) and *E. coli* FtsEX-EnvC (11) have revealed structural arrangements of the FtsEX heterodimer complex with a single extracellular binding partner which are the subject of a recent review (25). In total we now have a collective understanding of FtsEX function as enabling a conformational change in the extracellular hydrolase protein, presumably to position the hydrolase active site within the PG target layer as well as structural elements adjacent to the hydrolase that control hydrolytic activity.

It is worth noting when considering the peptidoglycan layer in *S. pneumoniae* that it is made of direct and indirect cross links in which the side chain of Lys at the third position of the peptide stem can be modified with L-Ser-L-Ala or L-Ala-L-Ala (as a result of amino acyl-tRNA dependent MurM and MurN action (26–28)). Altered stem peptide content and the resulting increase in PBP-mediated indirect crosslinking is found in highly-penicillin-resistant strains (29). A recent report of the catalytic domain of non-essential VldE (Spr1875) shows it to be a zinc-dependent L,D-endopeptidase which hydrolyses the cross-linked stem peptide and is induced via the VicRK (WalRK) two-component system under stress conditions (30), as is PcsB (31,32). Unlike previously mentioned PG amidases, VldE can produce a form of PG in which the first L-alanine in the peptide stem is left attached to the glycan sugar backbone (the hydrolysed bond is between L-Ala and D-Glu on the non-amidated synthetic peptides tested) (30). The role of VldE, which cannot replace PcsB, is unknown, but may be stress related (30). The LytA protein of *S. pneumoniae*, which is an N-acetylmuramoyl-L-alanine amidase produced in the cytoplasm, produces a fully denuded glycan strand but appears to play a role in fratricidal stress-induced lysis rather than in cell division (33).

In this study, we explore the structure-function relationship between the *S. pneumoniae* FtsEX complex with PcsB using *in vitro* and *in vivo* approaches that seek to explore the biological events that control PG hydrolysis. We explore the substrate specificity of PcsB’s CHAP domain (PcsB_CHAP_) whilst elucidating the molecular target and product of PcsB’s activity, defining the chemical nature of the hydrolase-modified PG following PcsB CHAP domain action. Finally, we demonstrate a putative regulatory element of PcsB_CHAP_ which is FtsX-independent, providing additional regulation of the tightly controlled FtsEX-PcsB PG hydrolytic activity.

## Results

### Co-overexpression of *Spn* PcsB and *Spn* FtsEX rescues phenotypic defects observed in *E. coli* when *Spn* FtsEX is overexpressed in isolation

We began our investigations by seeking to understand how PcsB is activated during cell division. To that end, we built upon previous data in which an *E. coli ftsEX* knockout strain demonstrated poor growth and cell death under osmotic stress conditions (15,34–36). Whilst the direct role of these proteins in *E. coli* in relation to stress is still unclear, the phenotype can be used as a tool for complementation experiments. Using an *ftsEX* knockout strain of *E. coli* (MG1655 Δ*ftsEX*, Supp. Table 1), we tested the viability of cells on rich (LB) or stress (LB without NaCl, LB0N) solid media supplemented with 1 mM IPTG in a spot-titre assay (Supp. Fig. 1). Under osmotic stress, the Δ*ftsEX* strain supports minimal growth, and can be complemented by pneumococcal *ftsEX* expression from the gene cassette in our pET-DUET vector construct, (Supp. Table 2) by more than two orders of magnitude (Supp. Fig. 1B). The *E. coli* MG1655 strain has no λDE3 lysogen for the T7 RNA polymerase system so expression will be reliant on recognition of *E. coli* RNA polymerase of the plasmid promoter system at a low or basal level (37,38). However, IPTG was included in the media in our experiments to placate interference from native *E. coli* LacI, thereby minimising repression of the plasmid’s T7-lac system. Notably, our results support a shared role for the FtsEX protein complex across multiple organisms and indicates a functional role for pneumococcal FtsEX in an *E. coli* host environment under low-salt conditions.

Utilising heterologous expression in this way also allows mechanistic examination of point specific mutations in *S. pneumoniae* essential genes and their protein products (39) which would be otherwise difficult to examine. Given this, we next chose to express these proteins in an *E. coli* (DE3) heterologous overexpression, in which there is wild-type expression levels of *E. coli* FtsEX-EnvC-AmiAB. This also allows testing of the individual components of the pneumococcal FtsEX-PcsB complex in a way that would not be possible in a pneumococcal strain background. The *pcsB* gene was separately cloned into a pCDF-DUET vector downstream of a *pelB* leader sequence (to enable the protein to be directed into the *E. coli* periplasm) also under T7 promoter control (Supp. Table 2), to allow co-transformation and heterologous expression of FtsEX-PcsB as a functional complex or as individual components in *E. coli*. Preliminary experiments suggested overexpression of *Spn* FtsEX at late-log phase resulted in a large proportion of highly filamentous *E. coli* cells, with no defined division septa along their length.

To explore this in more detail, cells were synchronised using DL-serine hydroxamate (40) prior to protein expression and grown to late-log/stationary phase in order to visualise phenotypic effects due to the presence of the recombinant proteins, rather than the transient effects of protein overexpression *per se* (41–43). The filamentous phenotype was reproduced during *Spn* FtsEX overexpression under these conditions (Fig. 1B) but not with *Spn* PcsB in isolation (Supp. Fig. 2G). By contrast, when PcsB was expressed in addition to FtsEX, the cells reverted to a rod-shaped phenotype (Fig. 1C). This unexpected but dramatic observation is consistent with FtsEX-PcsB acting in a functional manner during *E. coli* growth and division, rather than a more general non-specific effect of protein expression. The presence of PcsB within the context of the FtsEX-PcsB complex is apparently able to rescue the cell-division defect phenotype observed when FtsEX is overexpressed in isolation. Moreover, PcsB acts upon a non-cognate peptidoglycan substrate in this scenario in which the second and third amino acids in the peptide stem peptide are altered from native D-*iso*Glutamine to D-Glutamate and from L-Lysine to DL-Diaminopimelate, respectively.

**Figure 1.**
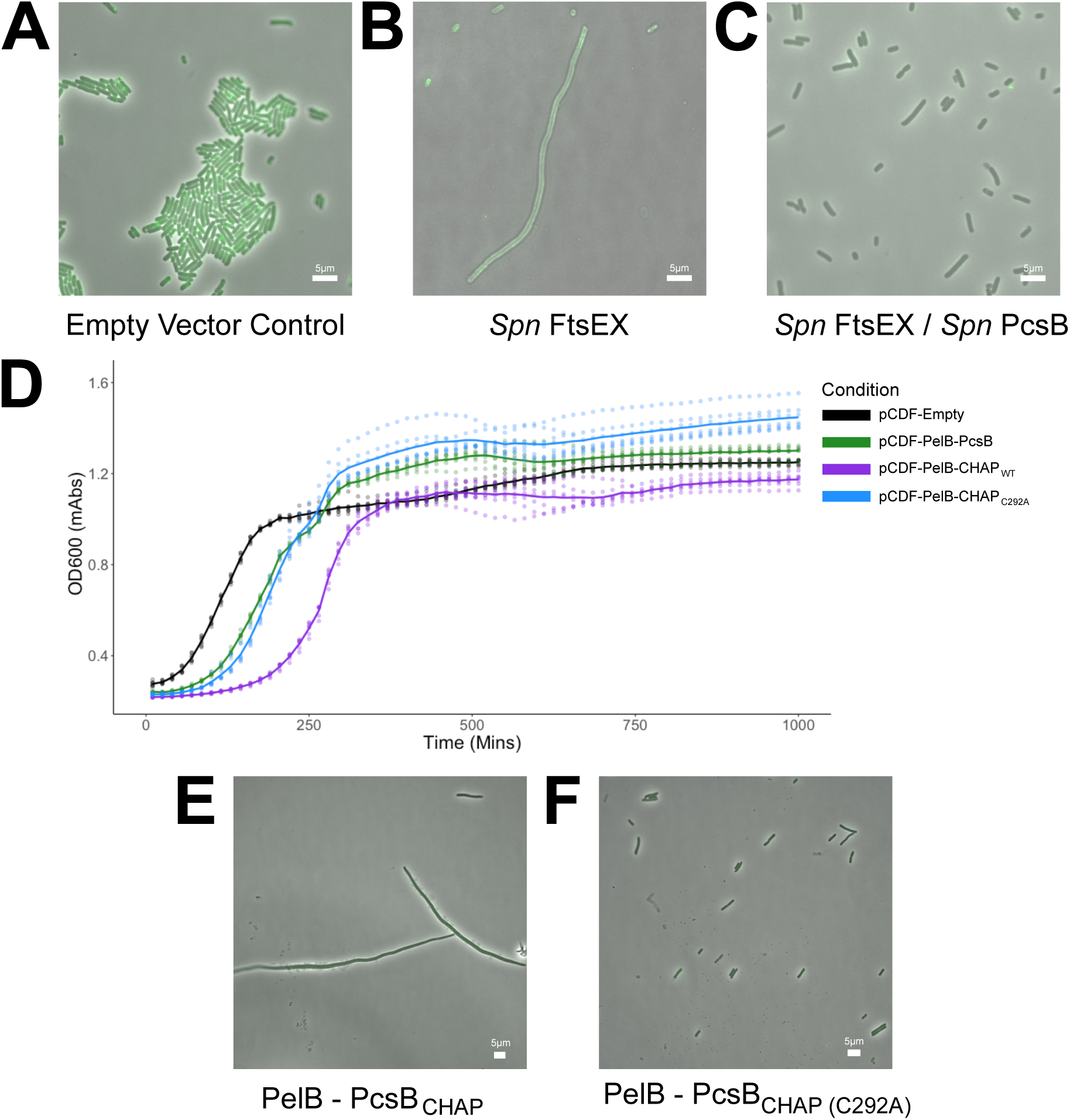
Co-overexpression of pneumococcal PcsB with pneumococcal FtsEX rescues the filamentous phenotype of FtsEX alone in *E. coli,* however overexpression of the PcsB CHAP domain in isolation forms filaments as well. A strain of BL21 (DE3) *E. coli* in which the parental *ftsZ* gene is tagged with *mNEONGreen* was used to visualise division septa during overexpression of pneumococcal proteins. Both empty vector (A) and FtsEX-PcsB (C) induced normal, rod-shaped *E. coli*, whereas expression of FtsEX alone induced filamentous cells (B). Growth curves of *E. coli* overexpressing empty vector, WT PcsB, PcsB CHAP domain or inactive PcsB CHAP (C292A) in the periplasm (PelB leader) (D) alongside representative microscopy images of active CHAP domain (E) or inactive CHAP domain (F) demonstrating cell division defects when the unregulated CHAP domain is active. Points mark OD reading at each time point; line marks, best fit line (mean of data). Magnification = 100x, scale bar (white) marks = 5 μm.

Subsequently, we sought to examine the structural elements required for the PG hydrolytic activity of PcsB, and specifically the CHAP domain of this protein (residues A274-D392, hereafter named PcsB_CHAP_) and assess what effect the active domain of PcsB has in isolation. PcsB_CHAP_ was cloned and expressed in a variety of constructs; in addition to the full-length wild-type and catalytically inactive C292A PcsB constructs, we made a truncated PcsB construct in which only the CHAP domain is directly preceded by a PelB leader sequence for periplasmic expression, thereby eliminating all proposed regulatory elements of the protein encompassed within the coiled-coil region (PcsB_CHAP_). Comparison of growth between *E. coli* overexpressing empty-vector, PcsB_CHAP_, PcsB_CHAP (C292A)_ and PcsB_WT_ cells showed a slight increase in the time taken to enter exponential phase for both the catalytically inactive PcsB_CHAP (C292A)_ and PcsB_WT_ compared to the empty-vector cells.

This effect was exacerbated with the PcsB_CHAP_ overexpression condition, indicating the active hydrolase has an aberrant effect on growth (Fig. 1 D). Phenotypic analysis of these cells revealed the PcsB_CHAP_ overexpression induces the same apparent phenotype as wild-type FtsEX, indicating the isolated hydrolase, lacking the proposed regulatory elements of the coiled-coil domain, induces cell division defects (Fig. 1 E and F). Taken together, these findings provide an effective visual reporter for activity and therefore function of the FtsEX-PcsB complex, which can subsequently be probed with mutations. In addition, an important insight into the activity of PcsB can be observed such that the CHAP domain of the protein shows sufficient *in-vivo* activity against a heterologous host’s peptidoglycan structure.

### Exploring the activation mechanism of PcsB via the FtsEX complex

In order to investigate these observations further and examine FtsEX-PcsB’s dependency on ATP turnover for activity of PcsB, we explored whether the cell-shape phenotype was dependent on the ATPase activity of FtsE within this complex, given the homology of the FtsEX-PcsB complex to ATP-dependent ABC transporters as well as the dependency for FtsE activity on the survival of pneumococcal cells (10). Mutations were made in the Walker A and B motifs of the *ftsE* gene to disrupt β-phosphate stabilisation (K43A) or Mg^2+^ binding (E165Q), as well as to the adenosine pi-stacking residue (Y13A); these mutants and FtsX were overexpressed in *E. coli* in the presence and absence of PcsB (Supp. Fig. 2). All three individual mutations resulted in the same filamentous phenotype as wild-type FtsEX when expressed in isolation or with PcsB (Supp. Table 5), implying that PcsB activation is driven by ATP turnover in the cytoplasm by FtsE. Nonfilamentous cells are also observed in these mutants, possibly due to compensatory mechanisms overcoming the deleterious effects that *Spn* FtsEX produces, or decreased induction of these proteins over time. Next, we examined if the PG hydrolytic activity of PcsB was also required to rescue the filamentous FtsEX-induced phenotype. An active-site Cys to Ala (C292A) mutant of *pcsB* was constructed and tested in the heterologous expression system (Supp. Fig. 2 and Supp. Table 5). Overexpression of PcsB_C292A_ in the presence of wild-type FtsEX did not rescue filamentation to rod-shaped cells (Supp. Fig. 2I and Supp. Table 5). Overexpression of either PcsB wild-type or C292A without *Spn* FtsEX has no obvious phenotypic effect in *E. coli* (Supp. Fig. 2 and Supp. Table 5).

Having confirmed this dependency on FtsE for PcsB activity, we then probed the activation mechanism of PcsB. It has been suggested in multiple FtsEX systems that a large-scale conformational change repositions the effector protein (PcsB here) thereby releasing the catalytic domain (13,18,24). To investigate this, we constructed a PcsB expression plasmid containing A194C and A231C mutations, which would link helix 2 and 3 of PcsB together under a direct disulphide (Fig 2A). We hypothesised that this would prevent the large-scale conformational change in PcsB driven by FtsEX from occurring, thereby preventing activity. We purified the FtsEX-PcsB complex in both wild-type form and with the PcsB (A194C/A231C) mutations (Supp. Fig. 3) for use in an ATPase activity assay. This assay uses a coupled enzyme system that monitors 7-methyl-6-thioguanosine (MESG) conversion to its free-base counterpart as a proxy for phosphate release and therefore ATP turnover activity (44). Using this system, we were able to show that FtsEX-PcsB_(A194C/A231C)_ has significantly reduced activity when compared to wild-type FtsEX-PcsB, but also that this activity can be restored by the addition of a reducing agent (DTT) into the system (Fig. 2B).

**Figure 2.**
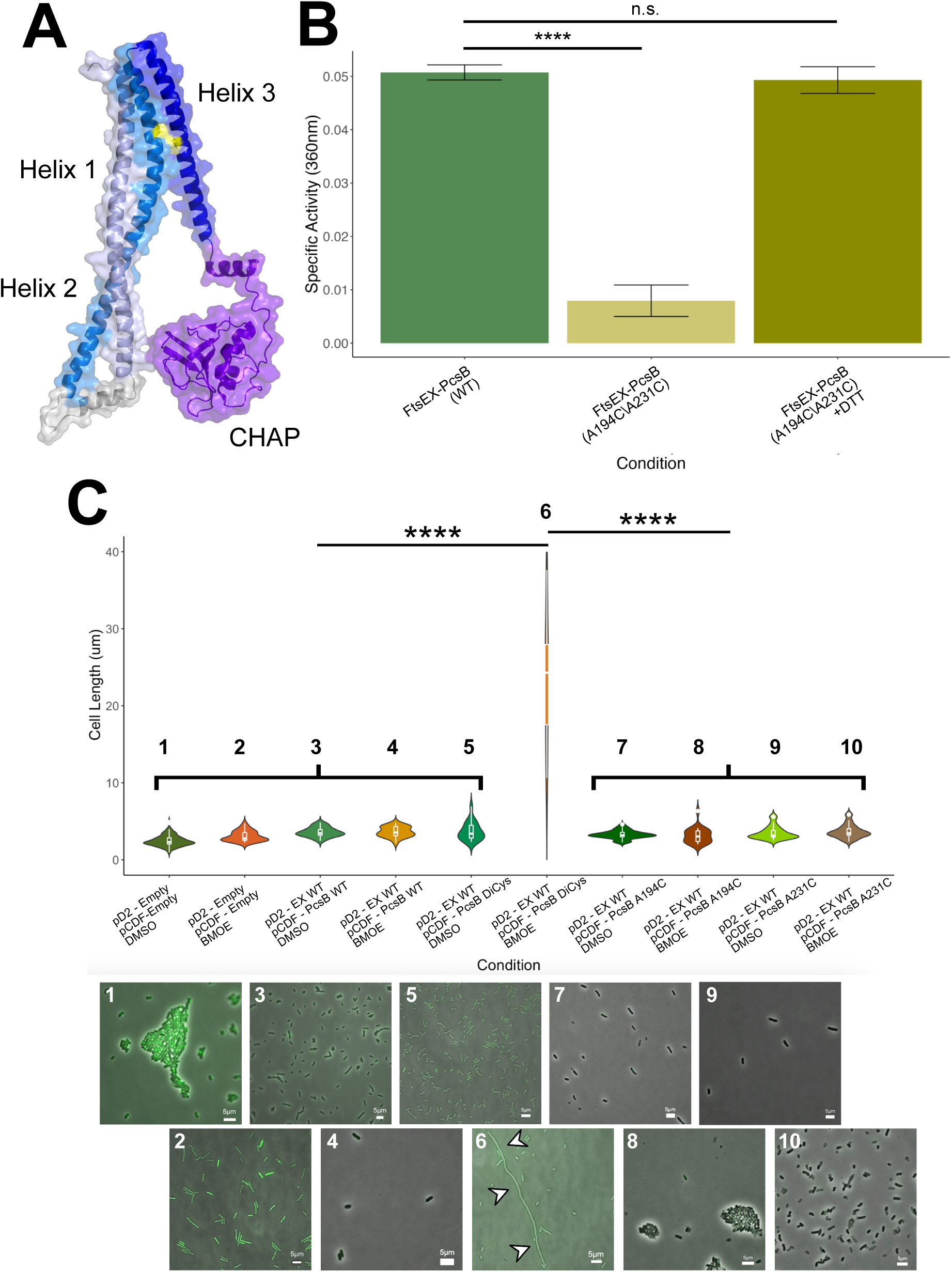
PcsB Activation Mechanism is Driven by Large-Scale Conformational Changes between Helix 2 and 3. Graphical depiction of PcsB structure highlighting Helix 1 (slate), 2 (marine) and 3 (blue) as well as the CHAP domain (purple) and the positions of A194/A231 (yellow) (A). A phosphate release assay demonstrates the requirement for mobility of PcsB Helix 3 for FtsE activity, revealing a novel feedback mechanism (B). Specific activity given in Abs.mins^-1^, error bars denote standard deviation spread. Heterologous overexpression of FtsEX-PcsB protein in *E. coli* demonstrating the specific restriction of conformation in PcsB that drives activation by Cys-based crosslinks; observed visually and via cell length measurements (C). Significance testing performed by t-test (n.s. – p>0.05, **** - p<0.0001).

This suggests a novel structural feedback loop present across the whole FtsEX-PcsB complex, whereby the conformation of PcsB drives the conformation of FtsEX as much as the traditional conception as the other way around. In exponentially growing *Spn* cells, a *pcsB*(A194C/A231C) mutant did not show a phenotype distinct from wild-type when treated with maleimide-based crosslinkers, possibly due to crosslinker interaction with other cysteine-containing proteins present on the pneumococcal cell surface, including the secreted form of PcsB which could be chelating the crosslinker. Therefore, to probe this feedback further, heterologous overexpression of this mutant as well as the mutations individually was carried out as above in the presence and absence of a maleimide-based crosslinker, bis-maleimidoethane (BMOE), to guarantee successful linkage of the Cys residues. All *E. coli* cells observed were the same length as wild-type, with exception of FtsEX-PcsB_(A194C/A231C)_ in the presence of the crosslinker BMOE (Fig. 2C), which showed significant filamentation indicative of a non-functional FtsEX-PcsB complex. This suggests the link is functioning as intended and is preventing activation of PcsB by inhibiting the conformational change. These experiments taken together reveal crucial insights into the mechanism of PcsB activation and the functional relationship between PcsB and the FtsEX complex.

### The CHAP domain of PcsB (PcsB_CHAP_) is an *iso*Glutaminyl-Lysyl D,L-endopeptidase

After uncovering aspects of the activation requirements for PcsB, we next moved to a mechanistic examination of the catalytically active portion of the protein, the CHAP domain of PcsB. We conducted a series of structure-function experiments in order to understand the peptidoglycan hydrolysis activity of PcsB at a molecular level. We expressed and purified PcsB_CHAP_ in order to explore its enzyme activity devoid of regulatory or inhibitory elements (Supp. Fig. 4). Initially we focused on using an industry-standard light-scattering assay designed originally for lysozyme (45). Continuous absorption monitoring at 450 nm confirmed that purified PcsB_CHAP_ was active, whereas the full-length PcsB protein was not (Supp. Fig. 5B and C). However, this assay relies on *Micrococcus luteus* cell wall as a substrate, which has fundamentally different biochemical properties to that of *S. pneumoniae* (e.g.: amidation state and cross-link composition) (46). To combat this, we chose to modify this assay to utilise whole peptidoglycan isolated from *S. pneumoniae* cells in which the enzymatic breakdown of the complex polymer can be monitored continuously by changes in absorbance at either 206 nm or 254 nm wavelengths to detect peptidase activity by carboxyl generation or glycan-strand cleavage, respectively (Supp. Fig. 5) (47). Lysozyme and recombinantly expressed pneumococcal LytA (Fig. 3A, B, Supp. Fig. 5 D and E) acted as controls for each detection wavelength due to their well-characterised endoglycosidase or endopeptidase activity, respectively (33,48,49).

**Figure 3.**
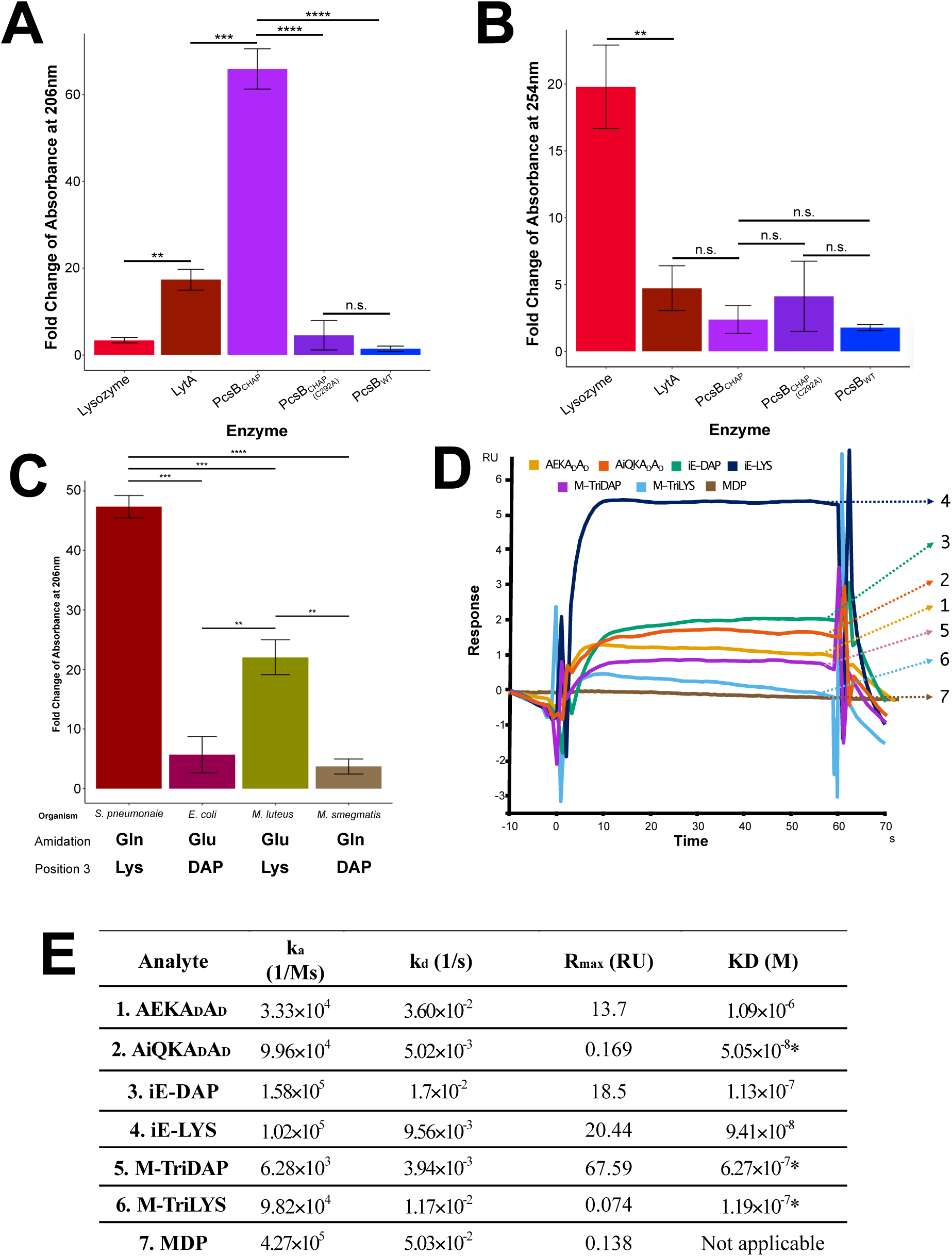
PcsB_CHAP_ displays endopeptidase activity with specificity for amidated, Lys-containing peptidoglycan substrates. Light-scattering-based assay using purified *Spn* peptidoglycan as a substrate measured at 206 nm (A) for carboxyl generation or 254 nm (B) for glycosidic bond cleavage. Light-scattering-based assay measured at 206 nm using PcsB_CHAP_ and respective peptidoglycan substrates (C). Overlayed SPR sensorgrams for general binding of substrates to PcsB_CHAP_ (D), summary of kinetic parameters derived from SPR analysis assuming 1:1 binding of PcsB_CHAP_ to various substrates (E). Single asterisks (*) in table E indicate significantly poor fittings as per Biacore Insight analysis software. Error bars denote standard deviation around mean generated from triplicate technical data. Significance determined by t-test. n.s. p>0.05, ** p<0.01, *** p<0.001, **** p<0.0001

Visualisation of the fold change of absorbance before and after the addition of protein (Fig. 3A and B, Supp. Fig. 5) demonstrated PcsB_CHAP_ displays comparatively high levels of activity at 206 nm (Fig. 3A), but minimal activity at 254 nm (Fig. 3B), consistent with hydrolytic activity upon the peptide stem rather than the glycan backbone. Recombinantly expressed and purified PcsB_WT_ and PcsB_CHAP (C292A)_ as well as natively expressed PcsB secreted and purified from pneumococcal cultures (Supp. Fig. 4) showed no significant activity at either wavelength (Fig. 3 A and B). This is consistent with the hypothesis that PcsB is only active within the context of the FtsEX-PcsB complex during division (22). We also extended this mutational screening to include the rest of the proposed active site (H343 and E360). Changing these residues to Ala resulted in total loss of activity for PcsB_CHAP (H343A)_ (Supp. Fig. 5); however, we found that PcsB_CHAP (E360A)_ could not be purified from the soluble fraction, and was poorly expressed even in the insoluble fraction. Based on interrogation of the PcsB structure (PDB: 4CGK) as well as our own structure(s), we concluded that E360 is likely involved in an sp2-hybridised pi-stacking network directly with N371, as well as with Y363 and F374. As a result, we were unable to confirm if PcsB’s active site functions with a catalytic diad or triad. However, we did confirm that these mutations which could be purified are mechanism specific by mutating N363, a residue near the catalytic triad to Ala, which showed no significant change in activity (Supp. Fig. 5).

From this, we aimed to further dissect the endopeptidase chemical specificity of PcsB_CHAP_ relative to the PG polymer in *S. pneumoniae*. We purified additional PG substrates from *E. coli*, *M. luteus*, and *Mycobacterium smegmatis*, providing the major variations in stem peptide combinations at positions two and three: D-*iso*Gln-Lys (*Spn*), D-Glu-diaminopimelic acid (DAP; *E. coli*), D-Glu-Lys (*M. luteus*), D-*iso*Gln-DAP (*M. smegmatis*) (50–52). Using these as PG substrates in the light scattering assay as described previously, we were able to ascertain that the D-*iso*Gln-Lys-containing substrate, as found in *S. pneumoniae*, was the preferred substrate over a D-Glu-Lys-containing substrate, as found in *M. luteus* (Fig. 3C). The DAP-containing PG stems found in *E. coli* and *M. smegmatis* were significantly poorer substrates in the assay, irrespective of the amidation of the second position amino acid, but notably still acted as substrates, as shown by phenotypic changes in overexpressing cells (Fig. 1E), though this effect is only seen over a much longertime frame. This indicates a wide substrate tolerance for the PcsB_CHAP_ active site as well as its potential to aid clearance of most Gram-positive cultures.

To probe the active site acceptance further, we used surface plasmon resonance (SPR) with selected commercially available peptidoglycan fragments to interrogate protein-ligand binding interactions further with the PcsB_CHAP_ protein bound to an SPR chip surface. Initial experiments were conducted with the active site knockout PcsB_CHAP (C292A)_, however no binding could be resolved with confidence. However, when the WT PcsB_CHAP_ protein was challenged with the pentapeptide stem peptide, amidation (of glutamate to *iso*glutamine) at the second position (A*i*QKA_D_A_D_) appears to induce an approximate 15-fold increase in binding KD compared to the non-amidated (AEKA_D_A_D_) isoform (Fig. 3E, in which the amidated peptide form corresponds to the species found in *S. pneumoniae* PG (Fig. 3C)). The performance characteristics of the pentapeptide in this experiment are non-ideal, however are consistent with on-chip hydrolysis of the pentapeptide by the PcsB_CHAP_ domain, and therefore allows for more in-depth analysis into the reaction kinetics, as well as pointing to a possible change in active site conformation upon substrate binding such that the active site Cys is involved in the interaction directly.

The dipeptides γ-D-Glu-mDAP (iE-DAP) or γ-D-Glu-Lys (iE-Lys) dissociate with highly similar affinities, so we extended this to include tripeptide species (classical NOD1 and NOD2 ligands (53–55)) that included the N-acetylmuramic acid residue. We therefore used MurNAc-Ala-D- *iso*Gln-Lys (M-TriLYS) and MurNAc-L-Ala-γ-D-Glu-mDAP (M-TriDAP) as ligands in the experiment and recorded slightly improved affinities (KD) compared to the dipeptide (Fig. 3 D/E). It should be noted that the presence of the amidation, as well as the sugars in M-TriDAP and M-TriLYS substrates, seems to be required for the enzyme recognition, as sensorgram profiles from the kinetics/affinity analysis of those substrates (Fig. 3E and Supp. Fig. 6) are consistent with on-chip enzyme activity, given the lower general resonance units (RU), which is proportional to the number of molecules bound to the surface, compared to other substrates, indicating that they are being actively hydrolysed and that the product is not remaining bound to the captured enzyme. The interaction with N-Acetylmuramyl-L-Alanyl-D-*iso*-Glutamine (MDP) gave no data that could be interpreted with our SPR experiment and unfortunately an N-Acetylmuramyl-pentapeptide species is not commercially available. Taken together, this binding analysis suggests that PcsB_CHAP_ substrate binding is dependent on both Lys and *iso*Gln presence, but may be enhanced by the sugar component of the ligand.

Finally, we determined the PG cleavage site upon which PcsB_CHAP_ acts. Incubation of synthetic pneumococcal stem peptide substrate L-Ala-D-*iso*Gln-L-Lys-D-Ala-D-Ala (A*i*QKA_D_A_D_) with PcsB_CHAP_ was initially separated using thin-layer chromatography (TLC) (Supp. Fig. 7A), which revealed a polar entity moving with the solvent front to a high position on the TLC plate that did not appear in any other lane. To investigate this further and validate the TLC observation, we therefore isolated the fragments of peptide and conducted mass spectrometry (MS) analysis on the products. The m/z peak corresponding to the full pentapeptide (488) was detected (Supp. Fig. 7B), which fragmented well under high energy and predictably such that the full pentapeptide could be assigned (Supp. Fig. 7C). Two other peaks were also identified at m/z values of 218 and 289 which are consistent with fragments for A*i*Q and KA_D_A_D_ originating from A*i*QKA_D_A_D_ peptide. High energy fragmentation of the 218 and 289 peaks allowed assignment of both peptide products respectively (Supp. Fig. 7D and E), leading to the conclusion that PcsB is an endopeptidase and specifically cleaves the bond between the *iso*Glutaminyl and Lysyl residues at position two and three in the PG stem respectively. This is further confirmed by our binding analysis of the MDP substrate to PcsB_CHAP_ (Fig. 3 D and E) which suggested low-to-no interaction, something expected of the product of the enzymatic cleavage.

To examine this further, we mutated the PcsB active site Cys292 to Ser in an attempt to trap the reaction intermediate in a more stable oxy-ester form compared to the native thio-ester. Following site-directed mutagenesis of the expression construct and subsequent expression and purification of the PcsB_CHAP C292S_ protein, the native pentapeptide substrate (A*i*QKA_D_A_D_) was incubated with the protein at an equimolar concentration. Following subsequent SDS-PAGE separation of the protein, an in-gel trypsin digest was carried out prior to mass spectrometry analysis of the peptide products. This revealed a shift in mass of +213 m/z, equivalent to the A*i*Q moiety bound via an ester to the now active-site Ser residue (Supp. Fig. 7F). This confirms both the scissile bond in this reaction as well as part of the mechanism of action of PcsB.

Whilst multiple attempts were made to crystalise PcsB_CHAP_ in the presence of substrate(s) or the oxy-ester intermediate, we were only able to capture the apo states of both PcsB_CHAP_ (PDB: 9R6W) and PcsB_CHAP C292A_ (PDB: 9R7G), (Supp. Fig. 8) which showed negligible changes in conformation when compared to the previously published structure (RMSD<0.3 Å) indicating that there is no change in the CHAP domain itself responsible for catalytic activation (22). By combining our in vitro activity data with our mass spectrometry data, we propose a mechanism for how the CHAP domain of PcsB catalyses the cleavage of the peptide stem within *S. pneumoniae* PG (Fig. 4).

**Figure 4.**
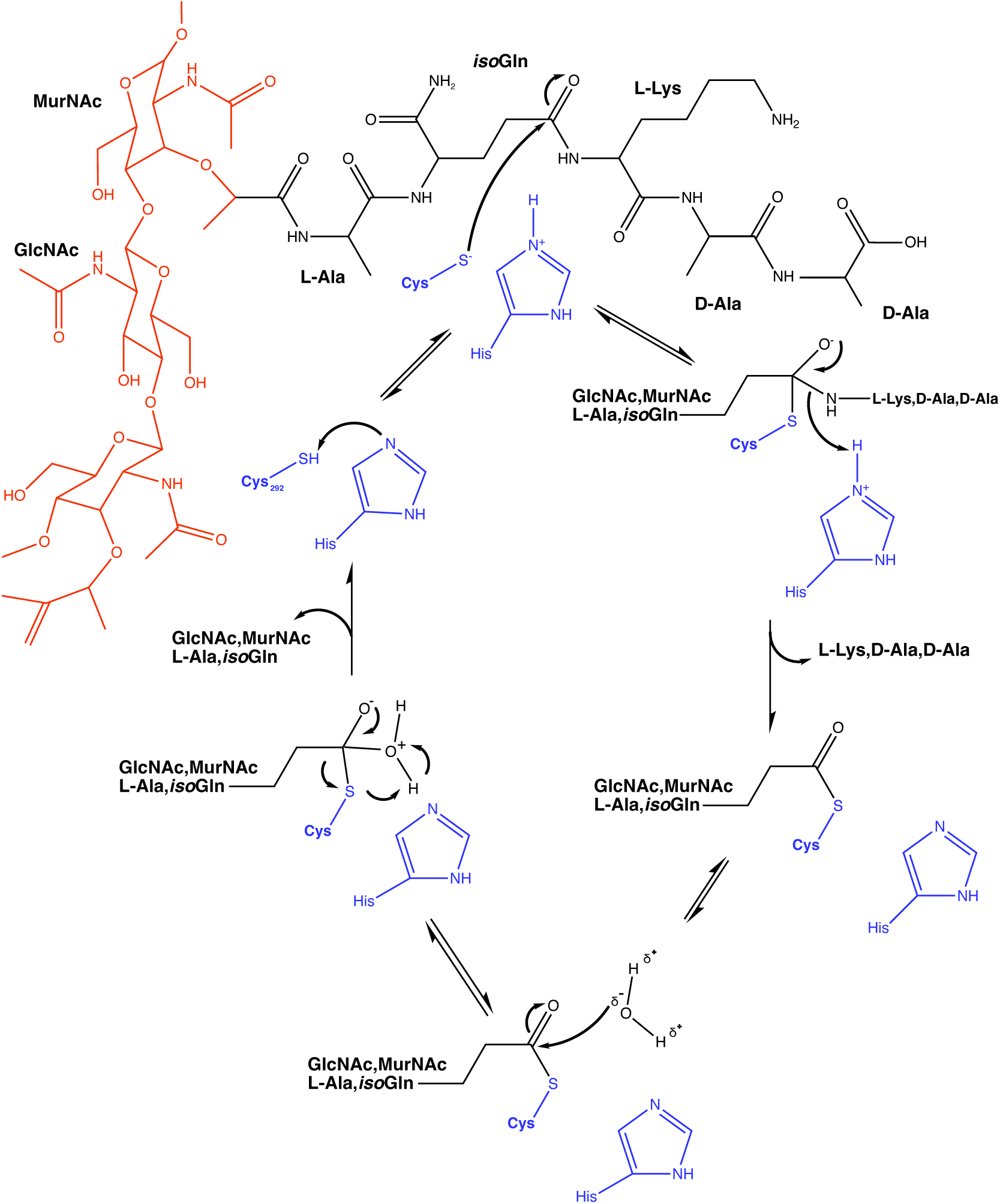
Proposed reaction schematic for PcsB. Blue moieties denote side chains belonging to PcsB protein structure (Cys_292_, His_343_), whilst red denotes the glycan strand and black denotes the pentapeptide side chain. In our proposed mechanism, Cys_292_, assisted by proton transfer to His_343_, initiates SN2 nucleophilic attack on the carbon atom of the peptide bond between *iso*Gln at position 2 and lysine at position 3 in the peptidoglycan side chain. The resulting tetrahedral transition state collapses, forming a PcsB-thioester intermediate, with expulsion of Lys-D-Ala-D-Ala, facilitated by protonation of the latter by His_343._ The thioester between PcsB-Cys292 and the remaining GlcNAc MurNAc L-Ala-*iso*Gln is subsequently hydrolysed via a second tetrahedral transition state (probably via a water nucleophile, consistent with a classical Cys protease mechanism (50)) to generate a carboxylic acid moiety on the peptidoglycan Ala-*iso*Gln stub and regenerate the free PcsB CHAP domain for subsequent rounds of catalysis.

Taken together, these results support a hypothesis that PcsB activity is optimised for (but not exclusive to) the pneumococcal PG substrate, rather than other forms of PG found in other bacterial species and that the enzymatic activity of the CHAP domain is consistent with an *iso*Glutaminyl-Lysyl D,L-endopeptidase, i.e. cutting between the second and third positions in the pentapeptide stem. Whilst PcsB is secreted into the media during pneumococcal growth (22), the lack of activation of the CHAP domain data (Fig. 3 (22)) argues against it having activity against competing bacterial species (i.e.: acting as a toxin). The ability of the CHAP domain to perform catalysis on a non-cognate substrate including the *E. coli* peptidoglycan is consistent with our previous co-overexpression and rescue complementation experiments of *Spn* FtsEX-PcsB in *E. coli* (Fig. 1 B and C, Supp. Fig. 2 I) having phenotypic effects consistent with a role in cell division. Finally, the conservation of structure seen between the CHAP domain in isolation determined here compared to that seen in context of the full-length PcsB structure suggests that D,L-endopeptidase activity requires major conformational changes in the coiled-coil domain, not the CHAP domain.

### The helix adjacent to PcsB’s CHAP domain has a biological function in regulating PcsB endopeptidase activity

In *E. coli*, as well as other Gram-negative bacteria, a pair of Zn^2+^-dependent amidases (AmiA/B) are known to bind FtsEX-EnvC and denude glycan strands via hydrolysis of the MurNAc-Ala ester bond (56,57). These amidases have also been shown to be regulated via helical structures adjacent to the hydrolytic domain of the proteins (20). Similarly, the *B. subtilis* endopeptidase, LytE, is regulated via a flexible loop from IseA that occludes the active site (21). A previous structure of PcsB (22) showed a similar short helix (N256-S267) close to the CHAP domain, followed by a flexible loop region (A269-T282). We therefore investigated if this region had a similar regulatory function within the PcsB system.

The full length *pcsB* gene was truncated to remove the coiled-coil domain, leaving only the 11-residue helix of interest, disordered region, and CHAP domain referred to thus as PcsB_CHAP+_ (residues T255-D392) (Fig. 5A). This was cloned into and expressed from pET-28b (Supp. Table 2), with the protein subsequently purified by Ni^2+^ IMAC (Supp. Fig. 4F). Assessment of this protein’s activity using the 206 nm light-scattering assay revealed no significant difference in activity from the full-length PcsB protein (Fig. 5B), in contrast to the isolated CHAP domain. This is consistent with a specific role for this helical-disordered region in inhibiting the catalytic activity of the CHAP domain.

**Figure 5.**
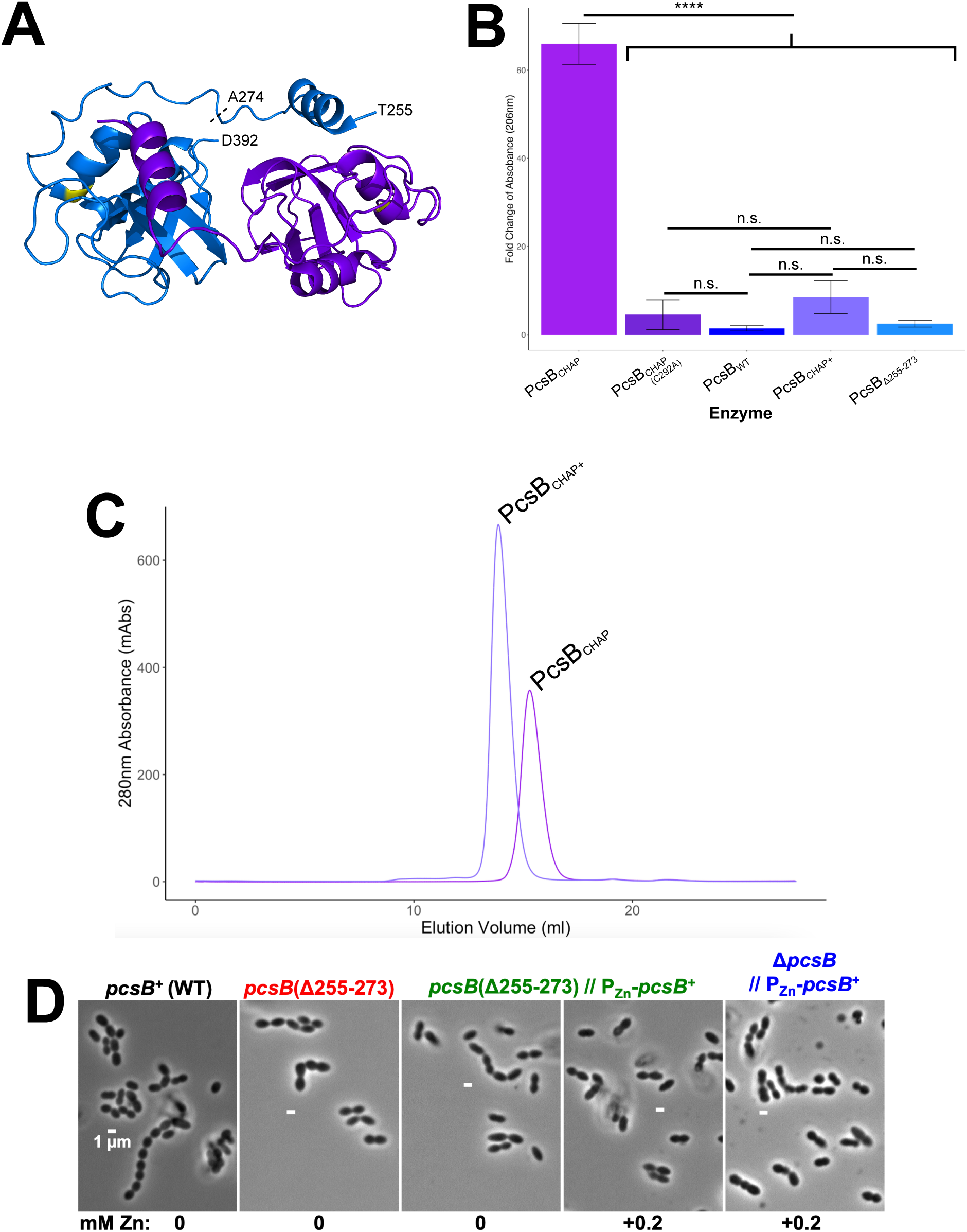
Short helical region adjacent to CHAP domain of PcsB appears to regulate activity of the PG hydrolase. Visual schematic from published PcsB structure (PDB: 4CGK) indicating CHAP domain with regulatory helix (PcsB_CHAP+_: residues T255-D392). Residue A274 indicates the start of the active endopeptidase CHAP domain, the proposed regulatory helix is shown between T255 and S265 (A). Blue and purple indicate two monomers, active-site Cysteine (C292) shown in yellow. Light-scatter-based assay measured at 206 nm (B) demonstrating lack of activity of the PcsB_CHAP+_ purified protein. SEC trace of purified PcsB_CHAP_ and PcsB_CHAP+_ proteins (C) indicating migration as a single species with no obvious aggregation. Representative phase contrast micrographs of WT and mutant *Spn* strains (D).

We hypothesised that this helix was being repositioned by the disordered region proceeding it such that the active site of PcsB was occluded. An alternative explanation for this observation is that the purified protein is aggregating via the helix and thus held in an inactive state in solution. We investigated this using size-exclusion chromatography (SEC), which revealed both the PcsB_CHAP_ and PcsB_CHAP+_ proteins migrate as equivalent monodispersed species (Fig. 5C). Neither protein showed evidence of aggregation, from which we suggest that regulation by the helix is most likely to be via direct interaction with the CHAP domain. It should be noted however that full-length PcsB missing only this helix (PcsB_Δ255-273_) is not catalytically active (Fig. 5B), which could suggest the coiled-coil domain inhibits the CHAP domain directly, or at least that the regulation does not involve this helical region.

Since the helix adjacent to PcsB’s CHAP domain (amino acids 255-273) largely abolished the *in vitro* endopeptidase activity of PcsB_CHAP_ (Fig. 5B), we asked if this helix regulated PcsB function within the pneumococcus. A pneumococcal mutant lacking this helix, *pcsB*(Δ255-273), expressed PcsB(Δ255-273) at levels comparable to WT (Supp. Fig. 9), but showed no growth phenotype in C+Y media (Supp. Fig. 10A).

Mutant *pcsB*(Δ255-273) cells were longer than WT, with increased cell volumes, and complementation with *pcsB*^+^ restored cells to WT size (Fig. 5D and Supp. Fig. 10). This increase in cell size was small but reproducible, and suggests that PcsB endopeptidase activity may be increased in the *pcsB*(Δ255-273) mutant *in vivo*, consistent with the idea that the 255-273 helix may partially inhibit PcsB function. However, the *pcsB*(Δ255-273) mutant was not more sensitive than WT to antibiotics such as penicillin, fosfomycin, vancomycin, ceftriaxone and cefuroxime, though it was slightly more sensitive to cefotaxime (Supp. Fig. 11). Taken together, these results suggest that the 255-273 helix of PcsB has a role in regulating PcsB endopeptidase activity in the pneumococcus, in addition to the ATP hydrolysis action of FtsE, transmitted through FtsX at the membrane and through the coiled-coil region of PcsB to the CHAP domain. In addition, antibiotic screening suggests that PcsB function could be intrinsically linked to the function of PBP2x, and is possibly exacerbated with an overactive PcsB, despite there being only a minimal effect on cell morphology. This may be clinically exploitable for downstream therapeutic applications.

## Discussion

The role of the FtsEX-PcsB in the context of bacterial cell division has been the subject of increased interest for a number of years. Recent structural elucidation of key components of the wider FtsEX system from multiple organisms has provided further mechanistic understanding (13,18,24,35), but a number of key biological questions still remain to be answered. One question is how is the PG-degrading activity of this protein complex controlled in order to facilitate the required cell division activity? This is especially acute for *S. pneumoniae* since in contrast to many other organisms, the pneumococcal FtsEX-PcsB protein complex is essential for cell survival. Whilst the *E. coli ftsEX* genes are conditionally essential, the growth defect phenotype observed with a Δ*ftsEX* strain under low-salt conditions appears to be rescued by complementation with *Spn ftsEX* alone (Supp. Fig. 1). Whether this complementation allows recruitment of EnvC-AmiA/AmiB or some other aspect of the protein-protein interactions required for assembly of the *E. coli* divisome is uncertain however. Conversely, overexpression of *Spn* FtsEX induces a filamentous phenotype in *E. coli* that can only be rescued with co-overexpression of the cognate hydrolase *Spn* PcsB (Fig. 1), further confirming a functional relationship between these three proteins (10,12,13,18,39).

This conclusion is enforced by examination of mutations in the *pcsB* and *ftsE* genes that overexpress PcsB or FtsE variants lacking PG hydrolase or ATPase activity respectively (Fig. 2 4, and Supp. Table 5). These results are consistent with a scenario in which the entire *Spn* FtsEX-PcsB system is reconstituted, representing active proteins in both cytoplasmic and extracellular compartments enabling peptidoglycan cleavage. Our detailed mechanistic and structural experiments reinforce this point and provide evidence for endopeptidase activity of PcsB (Fig. 3 and 4), as well as provide some of the first biochemical evidence for the conformational changes that result from activation of PcsB by FtsE (Fig. 5). The ability of the CHAP domain to degrade the non-cognate and chemically distinct *E. coli* peptidoglycan, though at a decreased efficiency, further supports this conclusion.

This leads to a second question, why is FtsEX widely regarded as being a complex that is recruited early in the assembly of the divisome? The answer to this may lie in part in the elucidation of the enzymatic activities of the hydrolases in question. The *E. coli* AmiA/B are amidases that cleave at the sugar-peptide linkage of the peptidoglycan structure, whereas PcsB is a D,L-endopeptidase that leaves a dipeptide stub (Fig. 4). As recently indicated by work on the SPOR domain of *E. coli* FtsN and its role in binding to denuded glycan and subsequent activation of the FtsWI complex (58), the formation of a modified form of PG may be a crucial marker or stage in the activation of later events in cell division. In *S. pneumoniae*, this marker may be the sugar-dipeptide stub. Whilst the order of divisome assembly in pneumococcus has not been fully established (but is partly implied from the *E. coli* system (3)), FtsEX-PcsB has been shown to associate with the FtsZ septal ring at midcell of predivisional cells (39,59,60). However as division progresses, FtsX (and by extension FtsE and PcsB) remain at the outer, peripheral (elongasome) synthesis ring, and are therefore separated from the closing septal annulus (60). This localisation pattern previously suggested a role in splitting existing septal PG, rather than the daughter cells directly (39,60,61). Given the observation here that PcsB’s mechanistic product leaves a sugar-dipeptide stub on the PG, it is possible that a second function of PcsB is to create a signal or space for subsequent PG remodelling to knit together septal and peripheral PG after cell separation is complete. It should be noted however that activation of PG synthases during division in pneumococcus is currently not well understood and is an area of active investigation.

Clearly peptidoglycan needs to be modified to allow cell division but at the same time new material must be synthesised to enable daughter cell separation. These processes are linked but distinctive and intuitively these processes must be highly controlled to prevent inadvertent lysis. The action of the CHAP domain of PcsB in *S. pneumoniae* or AmiA/B in *E. coli* will without doubt weaken the peptidoglycan by removing peptide crosslinks between adjacent glycan strands as part of this process. Whilst our focus here has been largely on the role of the extracellular hydrolase, the wider role of FtsEX in protein-protein interactions and assembly has been noted previously including interaction with FtsA (14) and FtsK in *E. coli* (62). It has also been suggested that there is a requirement for interactions between FtsEX and PG synthesis regulators (63), an area that is under current study. However, it should be emphasised that FtsEX-PcsB only associates to septal machinery during early division (2,60) as discussed above. Our phenotypic experiments are consistent with the hypothesis that *Spn* FtsEX interacts with the *E. coli* divisome as described (or at least influences the proper activity of the complex until PcsB co-overexpression induces displacement of the *E. coli* homologs), implying a shared and retained evolutionary history between these bacteria at the level of the divisome. It therefore seems likely that in pneumococcus, FtsEX has a dual role in early divisome recruitment and assembly whilst its ATPase function drives the activation of the extracellular hydrolase activity (to mark the site of division with modified peptidoglycan).

This subsequently poses the question of the role of PcsB and how it is controlled? We propose on the basis of the evidence presented here and elsewhere that PcsB functions as an essential “space making enzyme” during the cell division process, providing the separation within the sacculus for insertion of nascent glycan strands required for separation at the septal annulus. Tight regulation of PcsB activity is required, and this is primarily brought about by conformational change of the coiled-coil region of PcsB as the major regulatory mechanism, resulting from ATP hydrolysis in FtsE transmitted through FtsX (add key refs). The current hypothesis is that the third helix in the sequence of the PcsB coiled-coil structure extends upwards towards the PG layer thereby allowing the CHAP domain to act at the PG layer. Similar conformational changes have been suggested for homologous systems (11) and as presented in our model (Fig. 6), which is supported by our crosslinking experiments and suggests a novel structural feedback loop between PcsB and FtsE. We hypothesise that the short disordered-helical region upstream of the CHAP domain, which was noted within the PcsB structure (22), plays an additional role in regulation of the CHAP domain during activation of PcsB by FtsEX, in a similar way to the action of IseA of CwlO in *B. subtilis* (21). Whilst the *in vitro* activity assays suggested that this short region oblates the activity of the CHAP domain completely (Fig. 5), *in vivo* deletion of this region did not produce a constitutively activated protein. Instead, the cells showed a reproducible enlargement to their length and cell volume, implying that the major control mechanism is still through PcsB’s interaction with FtsEX, but that this short helix is likely to play a more subtle role in a CHAP-domain-specific manner.

**Figure 6.**
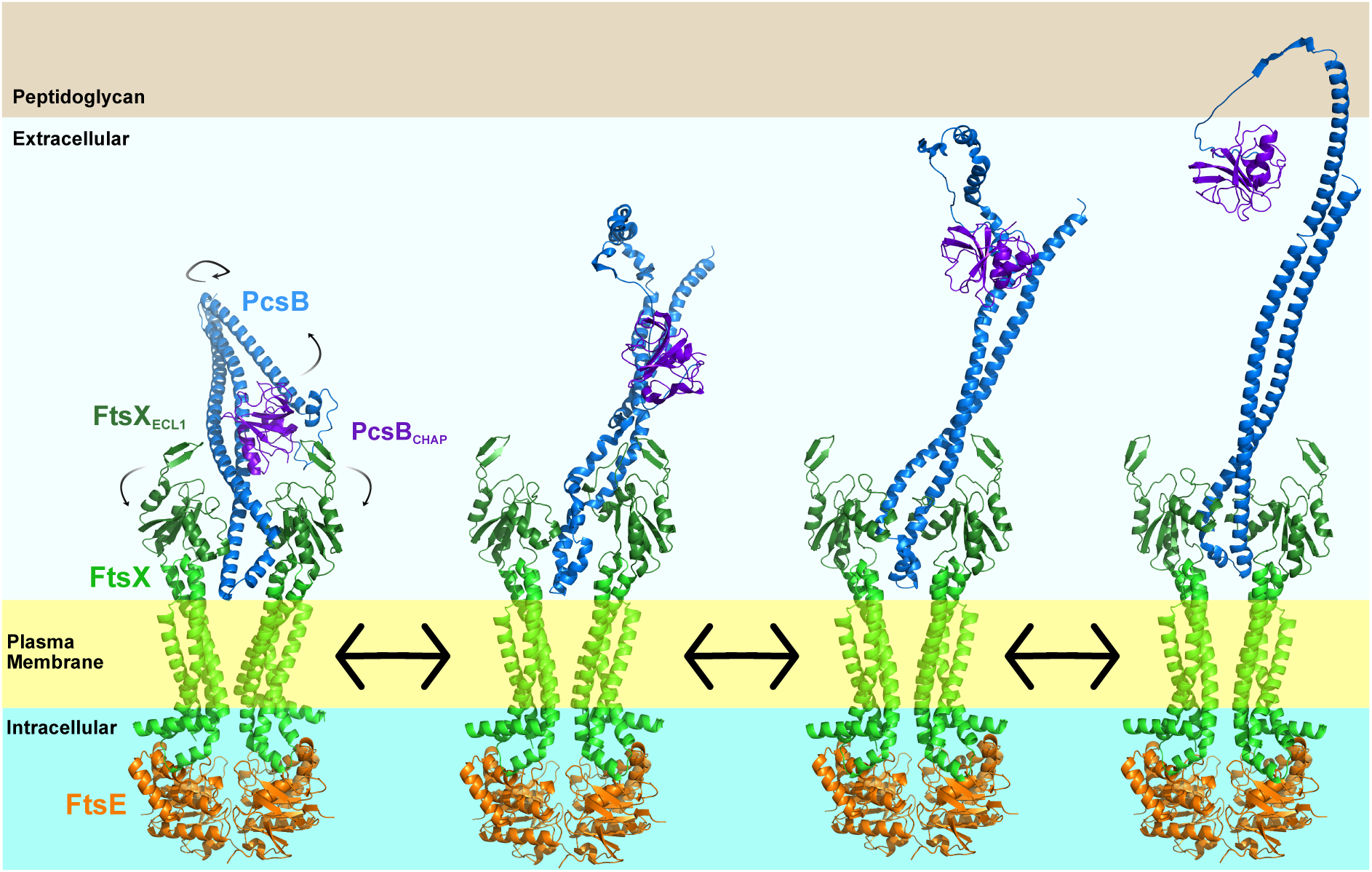
Proposed activation mechanism for PcsB via FtsEX. Turnover of ATP by FtsE in the cytoplasm induces a conformational change in FtsX embedded in the cytoplasmic membrane that in turn enables a conformational change in the coiled-coil region of PcsB. Extension of the coiled-coil region of PcsB liberates the CHAP domain from an inactive to an active state, allowing it to act on the peptidoglycan layer as an *iso*Glutaminyl-Lysyl D,L-Endopeptidase, producing a modified glycan with a dipeptide stub marking the site for further cell division activity. A short helical region adjacent to the CHAP domain provides additional regulation of endopeptidase activity. Models generated using AlphaFold3, intermediate steps generated using Pymol3 Morphing function.

Finally, we should ask the question of why *Spn pcsB* is essential? Taking the data presented here and elsewhere on balance, we propose that *Spn* PcsB has an essential role in *S. pneumoniae*, which classically has been in relation to cell separation (22,64). Here we introduce an additional role to that of septal annulus ring splitting previously proposed (60,61) as the protein appears to produce a modified form of peptidoglycan which marks the area of the cell required for synthesis of new material following division. This activity is controlled by ATPase activity in FtsE inducing mechanochemical confirmational change through FtsX and into PcsB to activate D,L-Endopeptidase activity. Additionally, FtsEX makes interactions with other parts of the divisome apparatus early in cell division as a coordinating element in the process which appears to be a conserved function across species. Whilst the enzymatic process of marking the site of division is performed by multiple systems in other bacteria including *E. coli* where the FtsEX-EnvC-AmiA/AmiB function can be compensated by multiple redundancies, in *S. pneumoniae* the role of FtsEX and PcsB is to make a MurNac-dipeptide stub within the peptidoglycan layer, marking the site of division, contributing to why PcsB is essential. In *E. coli* the SPOR-domain-containing protein FtsN recognises fully denuded PG and orchestrates later events in cell division. *S. pneumoniae* has no SPOR-domain proteins and thus a question arises as to what protein or proteins perform that function. The answer to this question is an active part of the research carried out in our laboratories and others.

In summary, we have demonstrated here that PcsB’s activity is specifically that of an *iso*Glutaminyl-Lysyl-endopeptidase that acts against pneumococcal cell wall PG and is predominantly regulated via the coiled-coil region, but also by a short regulatory helix that acts specifically on the CHAP domain. This work provides further insight into the molecular events to activate PcsB via ATP hydrolysis in FtsE, enabling mechanochemical conformational change from inside the cytoplasm via FtsX to PcsB on the extracellular surface and new molecular level detail on the mechanism and chemical features of PG cleavage during pneumococcal cell division. These data may contribute to further exploration of PcsB as an Achilles’ heel to target for future selective antimicrobial strategies against pneumococci (39).

## Acknowledgements

SN was supported by MRC Doctoral Training Partnership grant MR/N014294/1 and a Medical and Life Sciences Research fund award. NB was supported by a Ph.D. studentship to the Midlands Integrative Biosciences Training Partnership BBSRC grant BB/J014532/1 and by the Howard Dalton Centre at the University of Warwick under its small grant scheme award to support this work. SC was supported by MRC Infection and Immunity award MR/Z504245/1. Research in the laboratory of DR is supported by MRC grants MR/Z504245/1 and BBSRC grant BB/Y003187/1 and previously by a sabbatical Schaefer Research Scholars Program Awards to Columbia University NYC. Research in the laboratory of MW is supported by NIH Grant R35GM131767.

## Materials & Methods

### Spot Titre Assay

*E. coli* parent and *ftsEX* knockout strains (Supplementary Table 1) were gifted by Jonathan Cook, (Crow Lab, University of Warwick) and subsequently transformed with pET-DUET1-FtsEX construct containing the pneumococcal *ftsEX* gene cassette (Supplementary Table 2). Cells were subsequently cultured in LB (10g/L NaCl) broth with appropriate antibiotic selection up to an OD_600_ of around 1 before being serially diluted in PBS and spot plated onto LB agar or LB0N (0g/L NaCl) agar plates (2x biological replicates). Plates were incubated at 20°C overnight and imaged using ChemiDoc Imaging system (BioRad).

### *E. coli* Growth and Microscopy

*E. coli* BL21 (DE3)*_ftsZ_*_-mNEON_ cells were transformed with the construct(s) of interest. For growth curve analysis, cells were diluted to OD_600_ = 0.05 from overnight cultures and added to Cyto-One 96-well plates (StarLabs) in 2YT-AI media with appropriate antibiotics in triplicate (technical repeats) from 2 separate overnights. The plate was then incubated at 37°C for 24hrs in a BioTek Cytation 5 plate reader (Agilent). OD_600_ readings were taken every 15mins over this period. All repeat data visualised as points with the mean visualised as a best-fit line using ggplot package via R (V2024.04.1+748). For microscopy, overnight cultures were diluted in fresh LB 1:1 before being incubated with 1mg/ml DL-Serine Hydroxymate for 90 min. Cells were washed in LB before being diluted to OD_600_ = 0.05, and cultured at 37°C with antibiotic selection in 2YT autoinduction media (Formentum Cat#: AIM2YT0210) until OD_600_ = 0.8-1.0. The temperature was then reduced to 20°C overnight to allow expression of proteins. The following morning, cells were placed onto microscopy slides using a 1% agarose pad and imaged on Leica DMi8 Inverted Microscope. Measurements were taken of around 100 cells per condition across multiple images. During crosslinking experiments, the maleimide-based crosslinker bis-maleimidoethane (BMOE, Thermo Fisher) was included in growth culture after the synchronisation at a final concentration of 5 mM.

### Protein Expression and Purification

For soluble proteins (PcsB_CHAP_), constructs were transformed into BL21 (DE3) Star:pRosetta *E. coli* and grown overnight at 37°C. Cultures were then diluted into 2YT growth media with appropriate antibiotic selection up to an OD_600_ = 0.4-0.6, at which point IPTG was added to 300 μM. The temperature was reduced to 28°C and induction proceeded for 3 h prior to cell harvest at 6000 x *g*. Cells were then resuspended in 50 mM Tris-HCl (pH 7.5), 500 mM NaCl, 10% Glycerol, 3 mg/L Lysozyme and sonicated to lyse the culture before cell debris was pelleted at 40,000 x *g*. The soluble fraction of the cells was then loaded on a FastFlow 5ml HisTrap column (Cytiva) that had been pre-charged with 50mM Tris-HCL (pH 7.5), 500mM NaCl, 10% Glycerol, 10mM Imidazole. Column washing proceeded using 20%, 50% and 100% (50mM Tris-HCL (pH 7.5), 500mM NaCl, 10% Glycerol, 500mM Imidazole) solutions, fractions of which were collected and analysed for protein content via SDS-PAGE. Fractions containing purified protein were buffer exchanged into 50mM Tris-HCL (pH7.5), 200mM NaCl, 10% Glycerol, flash frozen and stored at –80°C.

For PcsB purification from pneumococcal growth media, a *Spn* strain was created expressing PcsB-His6x (strain IU18999). See the “*Streptococcus pneumoniae* Strain Construction” methods section below, as well as Supplementary Table 3, for details. This strain was cultured statically in BHI ± 1% choline chloride overnight at 37 °C + 5% CO_2_. Pneumococcal cells were removed by centrifugation at 4000g before the media was loaded onto a FastFlow 5ml HisTrap column (Cytiva) that had been pre-charged with 50mM Tris-HCL (pH 7.5), 500mM NaCl, 10% Glycerol, 10mM Imidazole. Column washing proceeded using Tris-HCL (pH 7.5), 500mM NaCl, 10% Glycerol, 50mM Imidazole solution before elution using Tris-HCL (pH 7.5), 500mM NaCl, 10% Glycerol, 500mM Imidazole solution. Fractions containing purified protein were buffer exchanged into 50mM Tris-HCL (pH7.5), 200mM NaCl, 10% Glycerol, flash frozen and stored at –80°C.

For the membrane-bound complex FtsEX-PcsB, constructs were transformed into BL21 (DE3) Δ*acrB E. coli* and grown overnight at 37°C. Cultures were diluted into 2YT growth media with appropriate antibiotic selection up to an OD_600_ = 0.8-1.0, at which point IPTG was added to 300 μM. The temperature was reduced to 18°C and induction proceeded for 3 h prior to cell harvest at 6000 x *g* and subsequent storage at -20°C. The following day, cells were thawed and resuspended in 50mM Tris-HCl (pH 8), 137mM KCl, 5% Glycerol before being lysed by 3 passes through the high-pressure homogenisation system, EmulsiFlex-C3 (Avestin), at 25kpsi. Ultracentrifugation at 120,000g for 1hr was then used to isolate the membrane fraction which was resuspended in 50mM Tris-HCl (pH 8), 500mM KCl, 5% Glycerol, 2.5% SMALP. This was incubated with agitation at room temperature for 3hrs with protease inhibitors before the ultracentrifugation was repeated to remove the insoluble membranes. The supernatant containing the solubilised membrane protein(s) was then incubated overnight at 4°C with 2ml bed volume Ni^2+^-NTA resin, pre-charged with 50mM Tris-HCl (pH 8), 500mM KCl, 5% Glycerol, 10mM Imidazole.

The following morning, this suspension was flowed through a gravity column prior to washing with 50mM Tris-HCl (pH 8), 500mM KCl, 5% Glycerol containing 10, 20 or 50mM Imidazole before final elution in 500mM Imidazole. This was subsequently subjected to size-exclusion chromatography on a HiLoad Superdex 200PG (Cytiva) which had been pre-equilibrated with 50mM Tris-HCl (pH 8), 200mM NaCl. Fractions corresponding to peaks at 280nm were analysed by SDS-PAGE for the presence of bands for FtsE, FtsX and PcsB (25kDa, 32kDa and 45kDa respectively. Correct fractions were pooled and concentrated to 1-5mg/ml, aliquoted to 50μl, flash-frozen in liquid nitrogen and stored at -80°C.

### Light-Scattering-Based Peptidoglycan Degradation Assay

Cell wall PG was purified from overnight cultures grown on respective agar plates (*E. coli* – LB, *S. pneumoniae –* BHI (+5% defibrinated horse blood), *M. luteus* and *M. smegmatis* – Nutrient). Cells were collected via cotton swabs into vials containing 1M NaCl prior to being boiled for 1hr. Following centrifugation, and washing with sterile water, a DNase/RNase solution was added (15μg/ml and 60μg/ml respectively) and allowed to incubate. Further boiling the suspension(s) preceded centrifugation and sterile water washing before the final suspension as made up in 12.5mM NaPO_4_ (pH 5.3). This cell wall was then added to a UV-vis quartz cuvette to a final concentration of 0.04% in 25mM KPO_4_. Once a stable baseline at all wavelengths (206nm, 254nm and 450nm) was established using a Cary-200 Spectrophotometer, protein was added to a final concentration of 286μM and the initial rate recorded. Each protein’s activity was recorded in triplicate.

### Phosphate Release (ATPase Activity) Assay

7-Methyl-6-thioguanosine (MESG) (LGC Biosearch) was loaded into a 200 μl quartz cuvette at 400 μM final concentration in HPLC-grade water. This was used to zero the Cary 100 Spectrophotomer at 360 nm during incubation at 37°C prior to the addition of 5 mM ATP. This was allowed to reach a stable baseline before the addition of 100 nM final concentration of ATPase test protein (FtsEX-PcsB) to initiate the reaction. The rate of increase of absorbance at 360nm was measured as a proxy for enzyme activity, before the baseline rate was subtracted to give the specific activity. Measurements recorded in triplicate per protein, with the average plotted.

### Thin Layer Chromatography

Purified PcsB_CHAP_ was mixed with the synthetic pentapeptide A*i*QKA_D_A_D_ (Sigma) at a 1:5 molar ratio (69.2μM and 346μM respectively) in 25mM KPO_4_ and incubated at 37°C overnight. Samples were separated on aluminium-plated silica-60 TLC plates (Fisher) and spots dried before being separated using the mobile phase 3:1:1 n-butanol:acetic acid:water. Bands were visualised using ninhydrin stain for both technical replicates.

### Mass Spectrometry

PcsB_CHAP_ protein was incubated with synthetic pentapeptide A*i*QKA_D_A_D_ at a 1:5 protein:ligand molar equivalence overnight at 37°C as for TLC. Protein was removed by centrifugal filtration using a 3kDa cut-off filter, allowing the peptide fragments to pass through. These were collected and analysed by direct spray ionisation mass spectroscopy in collaboration with Cleidiane Zampronio at WPH Proteomics Facility (University of Warwick). Samples were electrosprayed directly (∼300 nL min-1) through a Triversa Nanomate nanospray source (Advion Biosciences, NY) into a Thermo Orbitrap Fusion mass spectrometer (Q-OT-qIT, ThermoFisher Scientific, Germany) in positive ion mode. The gas flow was set to 0.3 psi and the voltage 1.7 kV. Data were acquired from 150 to 500 m/z with 120 K resolution (at 200 m/z). The precursor ions 218m/z, 289 m/z and 488 m/z were isolated and submitted to HCD fragmentation and detected on the Orbitrap. MS/MS analysis was performed using collision energy 35. This method was repeated for PcsB_CHAP (C229S)_ reaction with A*i*QKA_D_A_D_, aside from the protein being digested with Tyrpsin prior to reverse phase C18 separation and subsequent electrospray.

### Surface Plasmon Resonance

Kinetic parameters for the interaction of PcsB_CHAP_ with AEKA_D_A_D_, A*i*QKA_D_A_D_ (Sigma), MDP, iE-LYS, iE-DAP, M-TriDAP & M-TriLYS (Invitrogen) were measured by multicycle Surface Plasmon Resonance (SPR) on a Biacore T200 (GE Healthcare) employing a NTA Series S sensor chip (Cytiva). Prior to the analysis, the chip was conditioned with 350 mM EDTA and activated at the start of each cycle with 0.5 mM NiCl_2_. Chip regeneration at the end of each cycle was achieved with 30 s injection of 350 mM EDTA. His-tagged PcsB_CHAP_ (ligand) was buffer exchanged into running buffer (0.22 µm filtered PBS + 1% DMSO) using Zeba Spin desalting columns (Thermo Fisher Scientific) and diluted to a final concentration of 100 nM. PG fragments used as analytes were resuspended in running buffer to a final concentration of 10 mg/ml and eventually diluted to the starting concentration of 125 µM. Upon ligand capture, the analytes were injected in decreasing concentrations, following a 1:2 serial dilution from 125 µM to 7.812 µM. An injection of 0 µM analyte was employed to be used as blank during kinetics fitting. Contact time for the interaction was kept at 120 s, and the complex was left to dissociate by injecting running buffer for 600 s prior to regeneration. The same overall procedure was used to obtain a sensorgram overlay of the general binding between the analytes and PcsB_CHAP_. In this method, 100 µM analyte was injected for 60 s onto the captured protein and let dissociate for 60 s. Measurements were taken across triplicate runs.

Data analysis was performed on the Biacore Insight Evaluation software. Sensorgrams from the general binding method were overlayed and normalized after the capture level was assessed to counter the baseline shifting of the captured ligand and to allow an easier amplitude measurement. Kinetic parameters (k_a_ = association constant, k_d_ = dissociation constant, R_max_ = maximum response signal, KD = equilibration dissociation) were obtained by employing a 1:1 Langmuir interaction fitting model to calculate the KD.

### Protein Crystallisation and Structural Determination

Proteins were purified as described above and concentrated to 8-10mg/ml. Simultaneously, 80 µL of each sparce-screen condition (Structure-screen, JCSG+, Morpheus, MIDAS or ProPlex from Molecular Dimensions) was added to the reservoir of an MRC 2-drop plate (SWISSCI). Using the Formulatrix NT8 robot, protein samples were mixed with each reservoir condition at either 1:1 or 1:2 (protein:condition) ratio by volume before being covered with an optically clear sealing sheet (Molecular Dimensions) and incubated at 18°C. Each drop was checked daily for the first 7 days, then once a week afterwards under an SZX16 microscope (Olympus).

Identified crystals (PcsB_CHAP_ grown in 100mM Sodium citrate(pH 5.6), 500mM NaCl, 4 % v/v Polyethyleneimine (Molecular Dimensions Structure Screen condi8on G7; PcsB_CHAP C292A_ grown in 100mM BICINE (pH 9), 10% w/v PEG 6000 (Molecular Dimensions JCSG+ conditon E10)) were loop fished into 25% ethylene glycol (made up in crystallisation condition) and flash frozen in liquid nitrogen for transport to Diamond Light Source in Uni-Pucks (Jena Bioscience). Data was collected on the i04 beamline, with a total of 3600 images taken across 360° of rotation and auto-processed using the xia3 pipeline before being re-processed using Aimless (65) to removed batches that correspond to crystal degradation. Following determination of monomer number using Matthews (66), and generation of molecular replacement template using Swiss-model (67–69), PhaserMR was used to generate a solution structure based on the diffraction data that could then be refined using Coot (70–72) & Refmac5 (73). Model quality was checked using in-built validation tools within Coot as well as the B-average programme (70–72).

### Size Exclusion Chromatography

Purified proteins were injected onto a Superdex 75 10/300 GL via an AKTA Pure system which had previously been washed with 1 column volume (CV) water and 1 CV 25mM Tris-HCL (pH 7.5), 200mM NaCl at appropriate flow rates. Flow was carried out against gravity and separation monitored at 280nm. Fractions were collected using an F9-C automated fractions collector prior to peak fractions being analysed via SDS-PAGE.

### Streptococcus pneumoniae Strain Construction

*Streptococcus pneumoniae* strains used in this study were unencapsulated derivates of serotype 2 strain D39W (74) and are listed in Supplementary Table 3. *Spn* genomic DNA and oligonucleotides listed in Supplementary Table 4 were used to perform overlapping fusion PCR and generate linear amplicons. For transformation, *Spn* cells grown in brain heart infusion (BHI, BD Bacto 237500) broth at 37°C (all *Spn* incubations were done in a 5% CO_2_ atmosphere without shaking) to an OD_620_ ≈0.030 were diluted 1:10 in BHI broth containing 10% (v/v) heat-treated horse serum, 10 mM glucose and 100 ng/mL synthetic competence-stimulating peptide 1 (CSP-1) (75). Cells were incubated at 37°C for 10 m to induce competence before addition of 30-100 ng amplicon DNA, then incubated an additional 1 h. 100-800 µL cells were then added to 3 mL melted NB soft agar (0.8% (w/v) Nutrient Broth (BD Difco, 234000) and 0.7% (w/v) Agar (BD Bacto, 214010)) containing selective antibiotics and plated on trypticase soy agar containing 5% (v/v) defibrinated sheep blood (TSA-BA, BD BBL, 221261) and incubated overnight at 37°C. Single colonies were isolated on two successive days to fresh TSA-BA plates with selective antibiotics and incubated overnight at 37°C. For storage, a single colony was grown at 37°C in 3 mL BHI broth with selective antibiotics to OD_620_ 0.1-0.15, mixed with glycerol (final 15% (v/v)) and stored at -80°C. For antibiotic selection, 0.3 µg/mL erythromycin, 250 µg/mL streptomycin and 250 µg/mL kanamycin were used. Where necessary, cells were supplemented throughout transformation, plating, single-colony isolation and storage with 0.4 mM ZnCl_2_/0.04 mM MnSO_4_ (Zn with one-tenth concentration Mn included to prevent Zn toxicity, referred to throughout as “Zn inducer”) (76) to induce gene expression through a zinc-inducible promoter at the ectopic *CEP* locus (77) in the pneumococcal chromosome.

### *Spn* Growth and Microscopy

To make overnight cultures, frozen glycerol stocks were inoculated into 4 mL BHI broth, serially diluted and incubated 12-16 h at 37°C. When needed, overnight cultures were supplemented with 0.4 mM Zn inducer. For growth curve analysis in C+Y media (78,79), 2 mL overnight cell culture in exponential phase were centrifuged at 21,100 x *g* for 5 m at room temperature (RT), washed in 1 mL C+Y media, centrifuged again, and resuspended in 4 mL fresh C+Y. Cultures were diluted to OD_620_ 0.003 in 5 mL fresh C+Y with or without 0.2 mM Zn inducer, incubated at 37°C, and OD_620_ monitored every 45 m.

For microscopy, 500 µL cultures at OD_620_ 0.1-0.15 were centrifuged at 21,100 x *g* for 3 m at RT. 470 µL of the supernatant was removed and the pellet was resuspended in the remaining liquid with gentle pipetting. 1.25 µL of resuspension was placed on a glass microscope slide and a glass coverslip was gently applied. A Nikon Eclipse E-400 microscope, Nikon NIS-Elements BR imaging software, and a Nikon DS-Qi2 camera were used to capture phase-contrast images with 20-50 ms exposure time. Lengths and widths of >100 cells from 2 biological replicates were measured using Nikon NIS-Elements BR imaging software. For each cell, aspect ratio was calculated as length divided by width, and cell volume was calculated as length times the square of the width. Cell volumes were then normalized to the median cell volume of the WT strain. Dimension parameters were compared between strains using a Kruskal-Wallis test with Dunn’s multiple comparisons test (GraphPad Prism v10).

### *Spn* Disc Diffusion Antibiotic Sensitivity Assay

Performed as described previously (80), with slight modifications. Briefly, frozen glycerol stocks were inoculated in 3 mL BHI broth and incubated at 37°C. When OD_620_ reached 0.1-0.15, cultures were diluted to OD_620_ 0.009 in fresh BHI broth. 50 µL of cells were added to 3 mL melted NB soft agar, poured onto TSA-BA plates and incubated for 15 m at RT to solidify. After incubation, a paper disc containing an antibiotic (BD Sensi-Disc^TM^, Penicillin 230918, Cefotaxime 231606, Tetracycline 230998, Fosfomycin 231709, Vancomycin 231034, Ceftriaxone 231634, Chloramphenicol 230733, Cefuroxime 231621) was placed in the center of each plate, and plates were incubated at 37°C. After 16 h, the diameter of the zone of inhibition was measured. Measurements were averaged from 3 biological replicates and compared to WT using an unpaired *t* test (GraphPad Prism v10).

### *Spn* Quantitative Western Blotting

Quantitative western blotting of *Spn* cell lysates was performed as described previously for anti-aPBP1a and anti-MpgA antibodies (81), with minor modifications. Briefly, overnight cultures were supplemented with 0.4 mM Zn inducer as needed, and overnight cultures in exponential phase were washed and resuspended as described previously and diluted to OD_620_ 0.003 in 5 mL fresh C+Y with or without 0.2 mM Zn inducer as needed.

Cell growth, cell lysis, SDS-PAGE, transfer to nitrocellulose, total protein stain and blocking were carried out as described previously. The primary antibody used was a rabbit anti-PcsB antibody (1:1,000 dilution), generated by ThermoFisher Scientific using full-length *Streptococcus pneumoniae* D39 PcsB (amino acid M1 to D392, with an additional N-terminal sequence of LEHHHHHH) expressed in *E. coli* and purified as described above. Secondary antibody blotting, washing, imaging and quantitation of protein amount was completed as described previously (81). Shown is a representative result from 2 biological replicates.

