## Supplementary Data for "Uncovering the Biological Function of the Peptidoglycan Hydrolase PcsB in Streptococcus pneumoniae"

**Supplementary Table 1 – List of *E. coli* strains used in this study**

| Organism | Strain | Description | Reference |
|---|---|---|---|
| ***E. coli*** | MG1655 | Parental strain for *ftsEX* knockout | Gifted by J. Cook (Crow Lab, University of Warwick) |
| | MG1655 *ΔftsEX::Kan* | Kan Cassette inserted in place of native ftsEX operon | Gifted by J. Cook (Crow Lab, University of Warwick) |
| | BL21 *ftsZ-mNEON Green* (DE3) | BL21 background in which parental *ftsZ* gene is fused to *mNEONGreen* | (82) |
| | BL21 Star p.Rosetta (DE3) | BL21 background containing *rne*131 mutation as well as Rosetta plasmid for tRNA complementation | New England Biolabs |

**Commented [KB1]:** Confirm that either MG1655 or BW25113 are used correctly here, in the main text and fig. legends

**Supplementary Table 2 – List of plasmids used in this study**

| Plasmid | Vector Background | Content | Tag |
|---|---|---|---|
| FtsEX | pET-DUET1 | *S. pn* (D39) *ftsEX* operon | N-term 6xHis (*ftsE*) C-term 6xHis (*ftsX*) |
| PcsB & $PcsB_{\Delta 255\text{-}273}$ | pCDF-DUET1 | *S. pn* (D39) *pcsB* preceded by PelB leader | C-term 6xHis |
| $PcsB_{CHAP}$, $PcsB_{CHAP\ (C292A)}$ & $PcsB_{CHAP+}$ | pET-28b | *S. pn* (D39) *pcsB* truncated to CHAP domain (274-end), | N-term 6xHis |

| | | | |
|---|---|---|---|
| | | CHAP (C292A) domain (27A-end) containing C292A mutation or CHAP+ domain (255-end) | |
| LytA | pET-28b | *S. pn* (D39) *lytA* | N-term 6xHis |

**Supplementary Table 3 – List of *S. pneumoniae* D39W strains used in this study**

| Strain number[a] | Genotype (construction)[b c] | Antibiotic resistance[d] | Reference or source |
|---|---|---|---|
| **IU1824** | D39 Δ*cps rpsL1* | Str$^R$ | (74) |
| IU4959 | D39 Δ*cps pcsB*-$l_0$-*FLAG*$^3$-$P_C$-*erm* | Erm$^R$ | (39) |
| IU5122 | D39 Δ*cps rpsL1* CEP::$P_C$-[*kan-rpsL*$^+$] | Kan$^R$ | (60) |
| IU13123 | D39 Δ*cps rpsL1 ezrA*$^+$ // CEP::$P_{Zn}$-*ezrA*$^+$ | Str$^R$ | (60) |
| IU14778 | D39 Δ*cps rpsL1* CEP::$P_{Zn}$-*pcsB* (IU5122 X CEP::$P_{Zn}$-*pcsB* fusion amplicon) | Str$^R$ | This study |
| **IU14806** | D39 Δ*cps rpsL1* Δ*pcsB*::$P_C$-[*kan-rpsL*$^+$] // CEP::$P_{Zn}$-*pcsB*$^+$ (IU14778 X Δ*pcsB*::$P_C$-[*kan-rpsL*$^+$] fusion amplicon) | Kan$^R$ | This study |
| **IU18999** | D39 Δ*cps rpsL1 pcsB*-$l_0$-$his_{6x}$-$P_C$-*erm* (IU1824 X *pcsB*-$l_0$-$his_{6x}$-$P_C$-*erm* fusion amplicon) | Str$^R$ Erm$^R$ | This study |
| **IU20370** | D39 Δ*cps rpsL1 pcsB*(Δ255-273) // CEP::$P_{Zn}$-*pcsB*$^+$ (IU14806 X *pcsB*(Δ255-273) fusion amplicon) | Str$^R$ | This study |
| IU20409 | D39 Δ*cps rpsL1 pcsB*(Δ255-273) // CEP::$P_C$-[*kan-rpsL*$^+$] (IU20370 X CEP::$P_C$-[*kan-rpsL*$^+$] from IU5122) | Kan$^R$ | This study |
| **IU20481** | D39 Δ*cps rpsL1 pcsB*(Δ255-273) (IU20409 X *CEP*$^+$ amplicon from gDNA) | Str$^R$ | This study |

[a]**Bold** strains are used in experiments; other strains are intermediates or used to make amplicons for strain construction.

[b]Amino-acid sequences of linker 0 ($l_0$): GSAGSAAGSG. Linker DNA sequence is codon-optimized for *Spn* as described previously (83).

[c]$P_c$-*erm* and $P_c$-[*kan-rpsL*$^+$] cassettes described previously (84).

[d]Antibiotic resistance markers: $Erm^R$, erythromycin; $Kan^R$, kanamycin; $Str^R$, streptomycin.

**Supplementary Table 4 – List of oligonucleotides used in this study**

| Primer | Sequence (5' to 3') | Template[a] | Amplicon Produced |
|---|---|---|---|
| **For construction of IU14778 (CEP::$P_{Zn}$-*pcsB*)** | | | |
| KW116 | CCGGTAGTGGGAAAACAACTATTGGTCGTGC | IU13123 | 5' CEP::$P_{Zn}$ with 32 bp of 5' *pcsB* |
| KB471 | AATAAAAGTGACGCTAAGATTTTTTTCTTCATTACATCGCTTCCTCTCTATCTTCCTTGT | | |
| KB472 | AGGAAGATAGAGAGGAAGCGATGTAATGAAGAAAAAAATCTTAGCGTCACTTTTATTAAG | D39 gDNA | $P_{Zn}$-*pcsB* |
| KB474 | TTTGTTACATATATTTATGCAGATTAGAAAGAACGCTGAGAAGGTGTTCTTTTTATATTG | | |
| KB473 | GAACACCTTCTCAGCGTTCTTTCTAATCTGCATAAATATATGTAACAAAACCTTCAGAAG | IU13123 | 3' *pcsB*-CEP with 27 bp of 3' *pcsB* |
| KW123 | GGCTTCTTGTTCAAATTTTCCCATTTGATTCTC | | |
| **For construction of IU14806 (Δ*pcsB*::$P_C$-[*kan-rpsL*$^+$])** | | | |
| CS237 | TGTTGGTGAAGTGGTTGCCACAACGCATAG | D39 gDNA | 5' Δ*pcsB* ::*kan-rpsL*$^+$ with 23 bp *kan-rpsL*$^+$ |
| BR266 | CATTAAAAATCAAACGGATCCTAAGCTACTTGAGAAACCATTACTGTACTTAATAAAAGT | | |
| Kan rpsL forward | TAGGATCCGTTTGATTTTTAATGGATAATG | $P_c$-[*kan-rpsL*$^+$] cassette | $P_c$-[*kan-rpsL*$^+$] cassette |
| Kan rpsL reverse | GGGCCCCTTTCCTTATGCTTTTG | | |
| BR231 | CTAAACGTCCAAAAGCATAAGGAAAGGGGCCCGGATGGTTCAATCCAACAACAACTTCTG | D39 gDNA | 3' Δ*pcsB* ::*kan-rpsL*$^+$ with 32 bp *kan-rpsL*$^+$ |
| CS238 | TCTGCTTCAACTGCTACTGCATCCTCACCT | | |
| **For construction of IU18999 (*pcsB*-$l_0$-$his_{6x}$-$P_C$-*erm*)** | | | |
| CS237 | TGTTGGTGAAGTGGTTGCCACAACGCATAG | IU4959 | 5' *pcsB*-$l_0$-$his_{6x}$ with 19 bp $P_C$-*erm* |
| KB612 | AAACAAATTTTGGGCCCGGTTAATGGTGATGGTGATGATGGCCAGAACCAGCAGCGGAGC | | |

| Primer | Sequence (5' to 3') | Template[a] | Amplicon Produced |
|---|---|---|---|
| KB613 | CTCCGCTGCTGGTTCTGGCCATCATCACCATCACCATTAACCGGGCCCAAAATTTGTTTG | IU4959 | 19 bp $I_0$-$his_{6x}$-$P_C$-*erm* 3' *pcsB* |
| CS238 | TCTGCTTCAACTGCTACTGCATCCTCACCT | | |
| **For construction of IU20370 (*pcsB*(Δ255-273))** | | | |
| CS237 | TGTTGGTGAAGTGGTTGCCACAACGCATAG | D39 gDNA | 5' *pcsB*(Δ255-274) |
| KB648 | AGCGTTTGTACTGTATGTTGGACGAACTTTTGCGTTTGCTGAAGCAAGTACTGATTGTTG | | |
| KB647 | AACAACAATCAGTACTTGCTTCAGCAAACGCAAAAGTTCGTCCAACATACAGTACAAACG | D39 gDNA | 3' *pcsB*(Δ255-274) |
| CS238 | TCTGCTTCAACTGCTACTGCATCCTCACCT | | |

[a]Genomic DNA of D39W was used as templates for PCR reactions, except for the $P_c$-[*kan-rpsL*$^+$] cassette (84).

**Supplementary Table 5 - Summary of microscopy findings, demonstrating that the recue phenotype is due to the FtsEX-PcsB complex being fully active.** Mutations in either the ATPase domain of *ftsE* (Y13A, K43A or E165Q) or the active site of *pcsB* (C292A) do not rescue the cells from forming filaments. N.P. denotes "not present", describing situations where only one plasmid is present within cells.

| FtsEX Amino Acid Changes | PcsB Amino Acid Changes | Effect | Phenotype |
|---|---|---|---|
| N.P. | N.P. | Parental Cells | Rods |
| WT | N.P. | Active FtsEX | Filaments |
| **WT** | WT | Active FtsEX – Active PcsB | Rods |
| **FtsE – Y13A** | N.P. | Inactive FtsE | Filaments |
| **FtsE – K43A** | N.P. | Inactive FtsE | Filaments |
| **FtsE – E165Q** | N.P. | Inactive FtsE | Filaments |
| **FtsE – Y13A** | WT | Inactive FtsE – no PcsB activation | Filaments |
| **FtsE – K43A** | WT | Inactive FtsE – no PcsB activation | Filaments |
| **FtsE – E165Q** | WT | Inactive FtsE – no PcsB activation | Filaments |
| **N.P.** | WT | PcsB present but inactive | Rods |
| **N.P.** | C292A | Inactive PcsB | Rods |
| **WT** | C292A | Inactive PcsB | Filaments |

**Commented [KB2]:** I tried to simplify the formatting. Hope that's OK

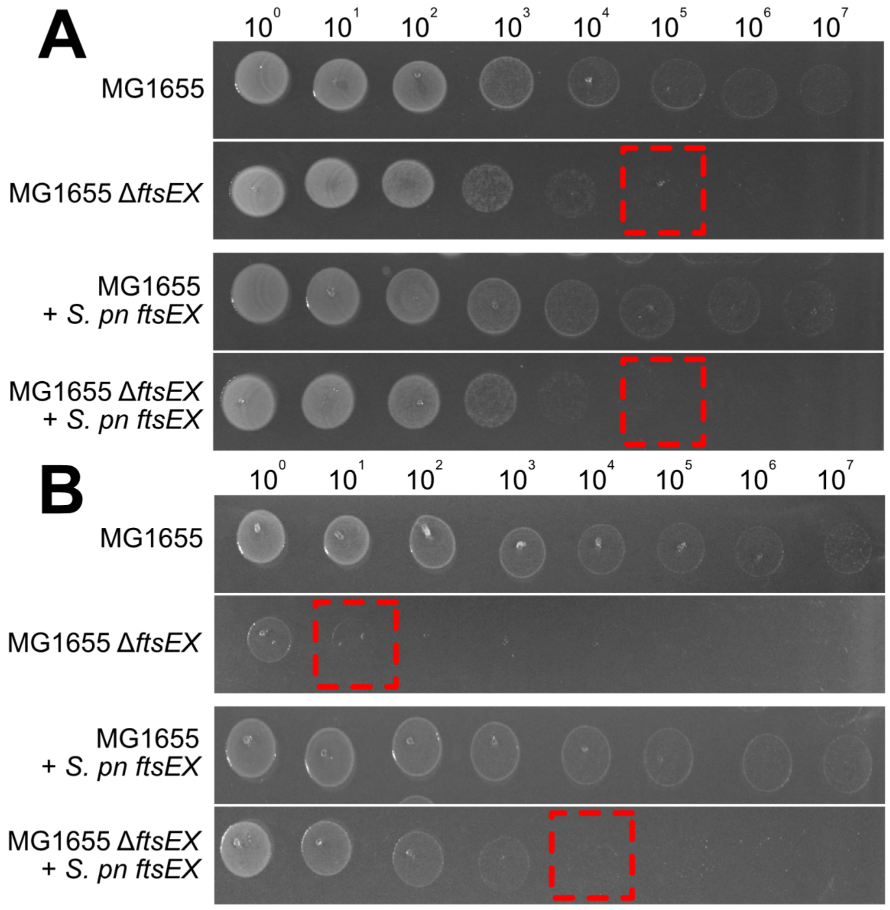


**Supplementary Figure 1 - Pneumococcal FtsEX complements the growth phenotype of *E. coli ΔftsEX.*** Spot-titre assay indicating cell survival at 10-fold dilutions from $OD_{600}$=1 culture ($10^0$). Cells were plated on osmoprotective LB (A) or hypoosmotic LB0N media (B), indicating pneumococcal FtsEX rescues growth by more than two 10-fold dilutions in low-salt conditions.

**Commented [KB3]:** Please include the number of biological replicates, either here or in the Methods.

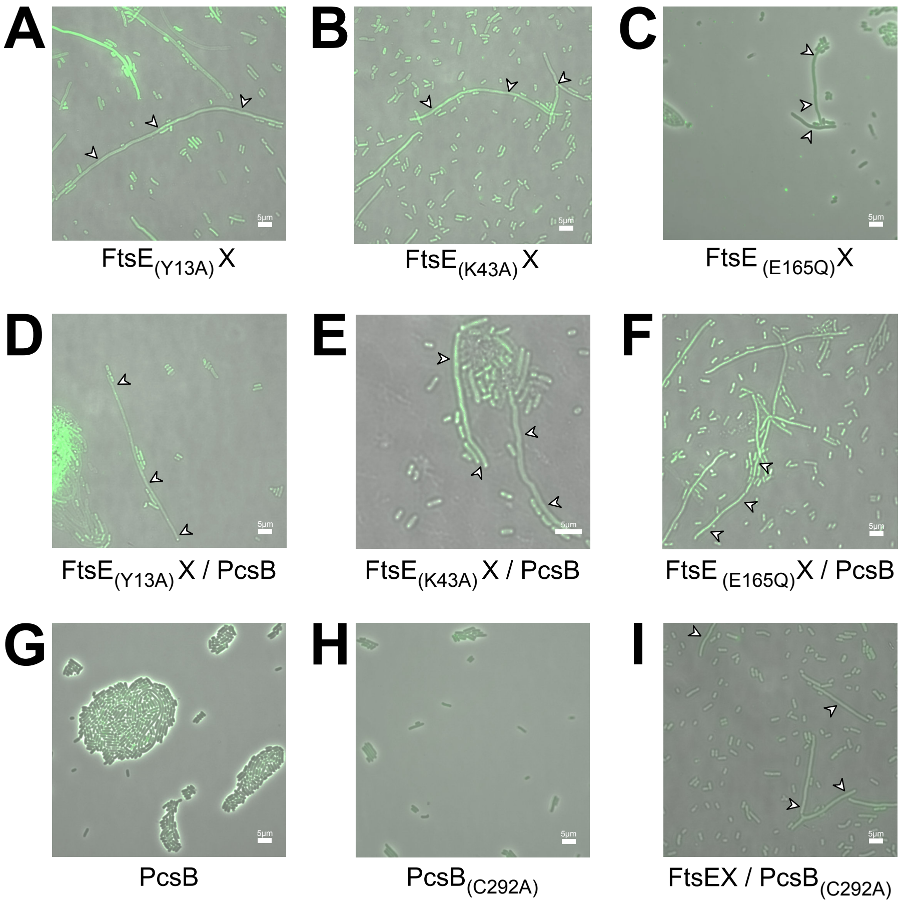


**Supplementary Figure 2 – Representative microscopy images demonstrating activity requirement for the rescue phenotype by FtsEX-PcsB.** Overexpression of proteins performed in a BL21 (DE3) FtsZ-mNEON background imaged after overnight induction (around 18hrs) in 2YT-AI media. White arrows indicate filamentous cells.

**Commented [KB4]:** The Methods state the overnight induction is in 2YTAI media. Please clarify

**Commented [KB5]:** Please include number of biological replicates, either here or in the Methods

**Commented [KB6]:** Remove panel B. For panel A: Y axis is cell length, not mean cell length. Include information on the number of biological replicates, how many cells were measured for each condition, what the whiskers of the box plot represent, the statistical tests used, and what p value **** represents

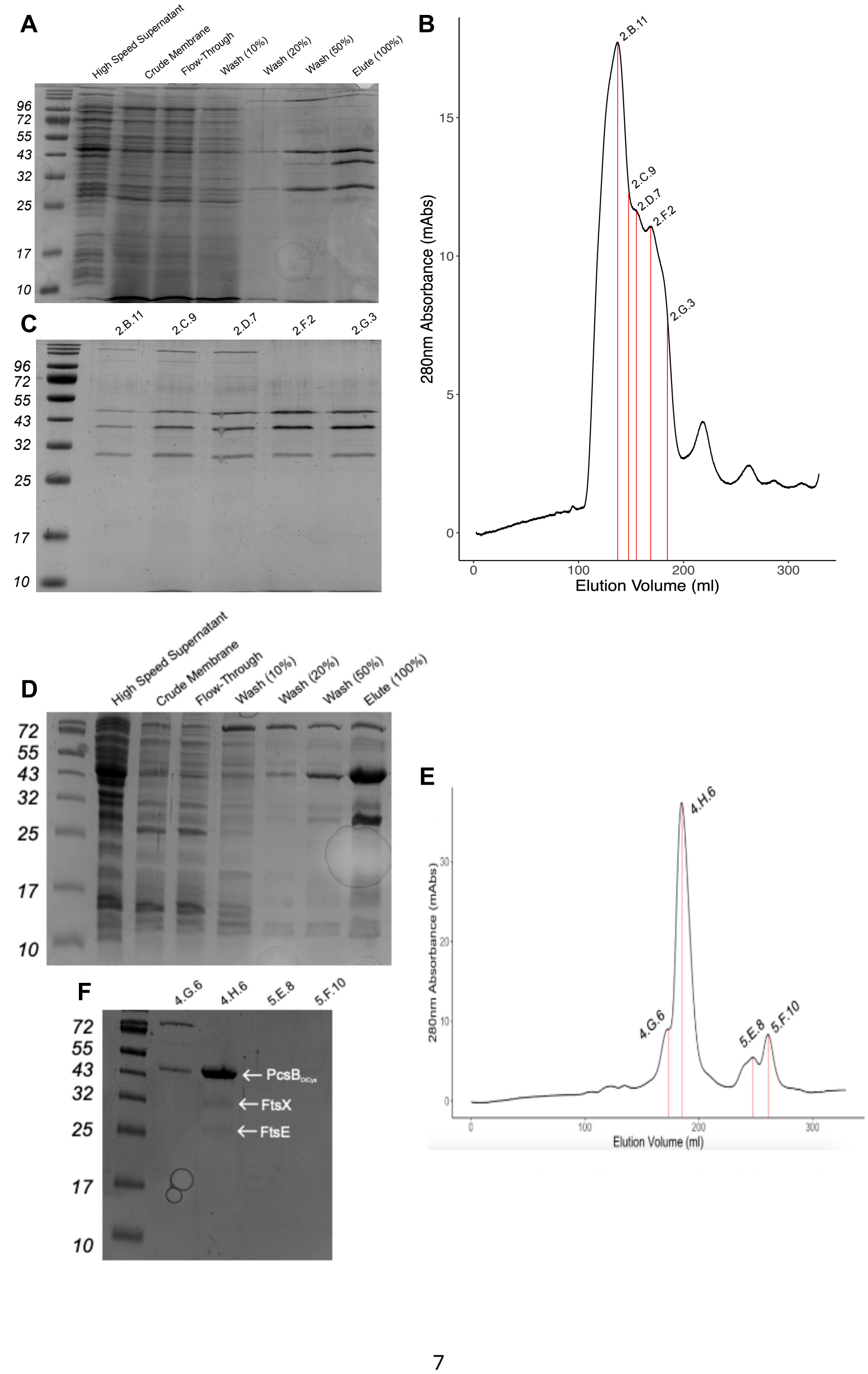
A
High Speed Supernatant
Crude Membrane
Flow-Through
Wash (10%)
Wash (20%)
Wash (50%)
Elute (100%)
96
72
55
43
32
25
17
10
B
2.B.11
2.C.9
2.D.7
2.F2
2.G.3
280nm Absorbance (mAbs)
Elution Volume (ml)
C
2.B.11
2.C.9
2.D.7
2.F2
2.G.3
D
High Speed Supernatant
Crude Membrane
Flow-Through
Wash (10%)
Wash (20%)
Wash (50%)
Elute (100%)
E
4.H.6
4.G.6
5.E.8
5.F.10
280nm Absorbance (mAbs)
Elution Volume (ml)
F
4.G.6
4.H.6
5.E.8
5.F.10
PcsB
FtsX
FtsE

**Supplementary Figure 3 – Purification of FtsEX-PcsB using SMALP nanodiscs.** Nickel IMAC purification of FtsEX-PcsB (A/D) alongside representative SEC trace (B/E) and subsequent SDS-PAGE analysis of peaks (C/F) for FtsEX-PcsB$_{WT}$ (A/B/C) or FtsEX-PcsB$_{(A194C/A231C)}$ (D/E/F).

**Commented [KB7]:** I know nothing about protein purification, but are you all confident with the amount of FtsE and FtsX purified with PcsB(A194C/A231C) in panels DEF? It seems low relative to the WT

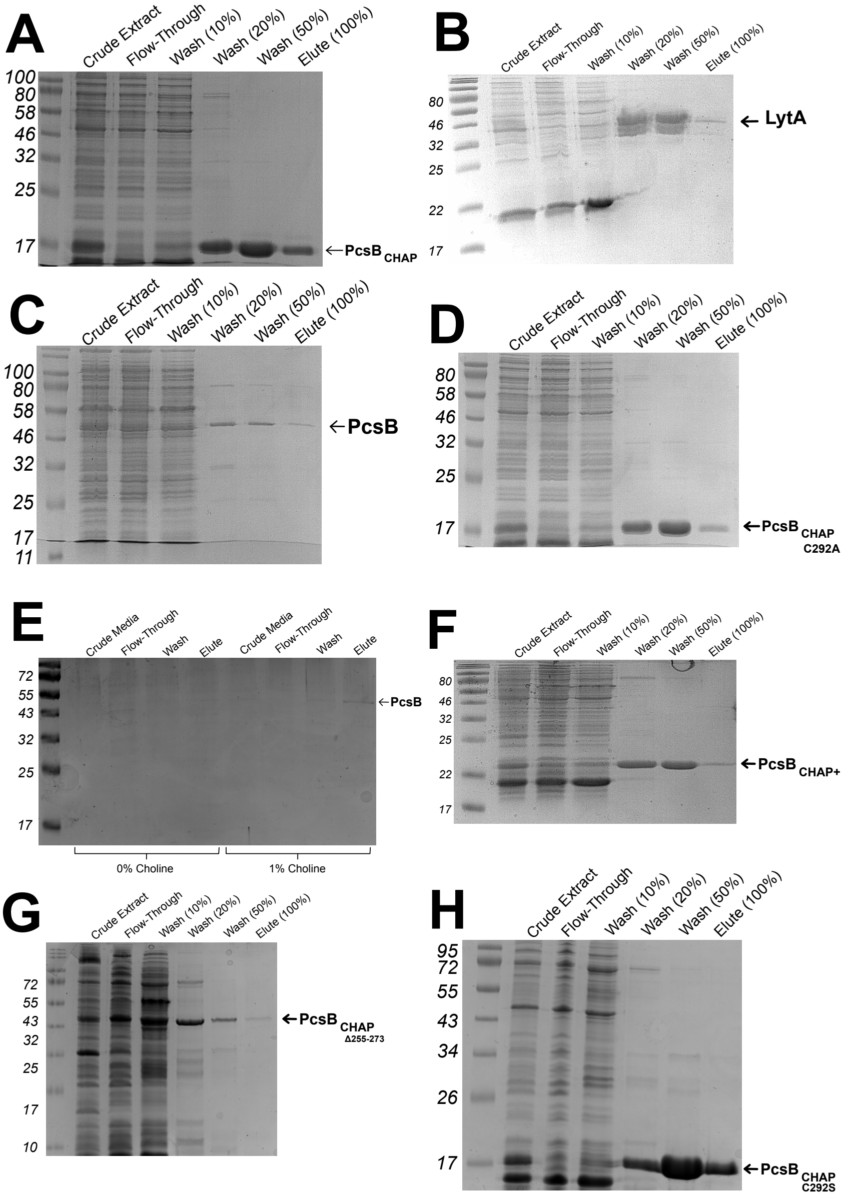


**Supplementary Figure 4 – $Ni^{2+}$ IMAC purifications of all PcsB proteins**. CHAP domain (A), Pneumococcal LytA (B), Recombinant wild-type PcsB (C), Catalytically inactive CHAP Domain (D), Pneumococcal PcsB purified from media (E), $PcsB_{CHAP}$ with helix T255-267 (F),

Recombinant wild-type PcsB with helix region 255-273 deleted (G), $PcsB_{CHAP\ (C292S)}$ mutant used for mass spec (H).

**Commented [KB8]:** Labeled 255-267 in the figure, please rectify

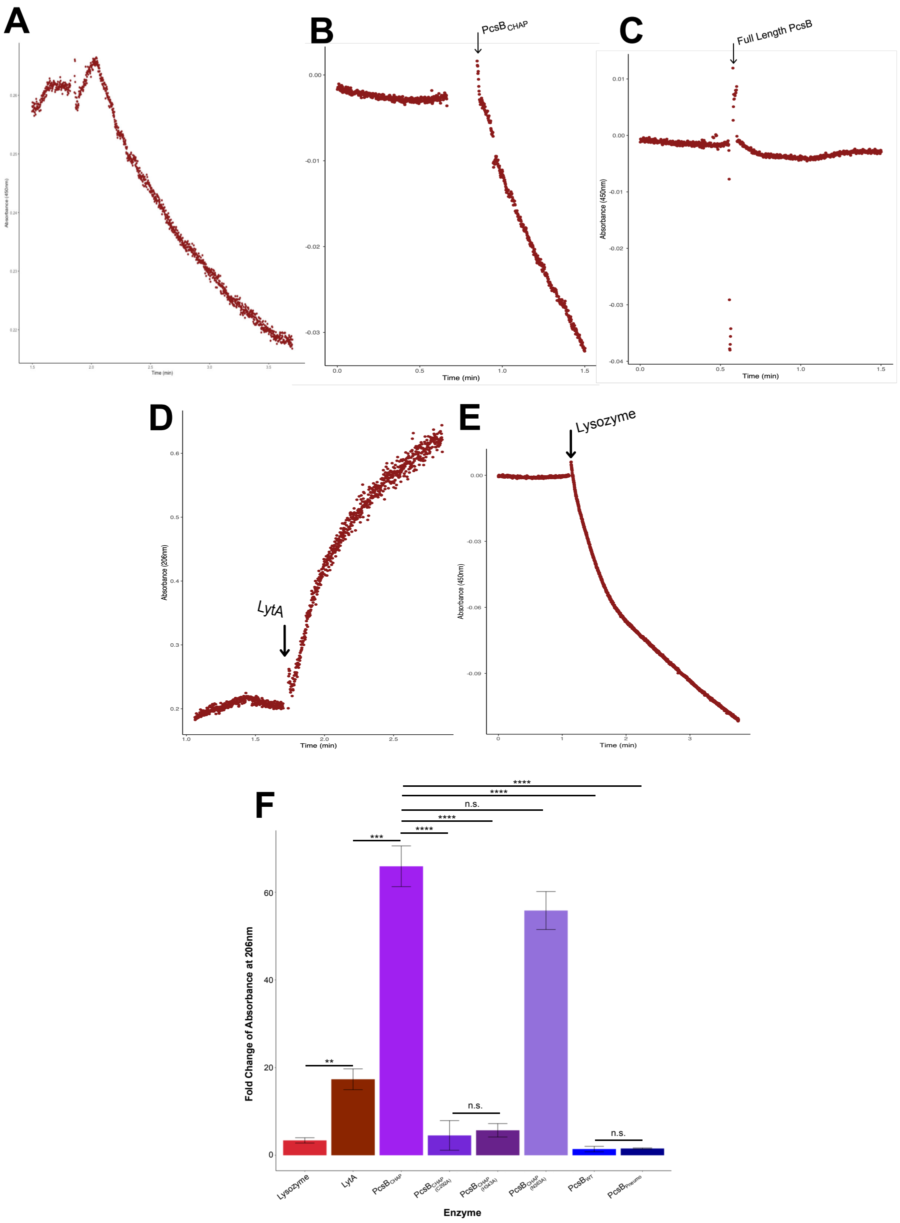

**Supplementary Figure 5 – Testing, Modification and Validation of 206/254 nm light-scattering assay.** Representative traces from commercially available 450 nm continuous absorbance-based light scatter assay measuring at 450 nm for Lysozyme (A), $PcsB_{CHAP}$ (B) and $PcsB_{FL}$ (C), where reduction in signal indicates activity. Modified assay using purified pneumococcal cell wall showing positive control signals measured at 206 nm with LytA (D) or 254 nm with lysozyme (E). Mean change in absorbance for all tested $PcsB_{CHAP}$ proteins in 206nm assay (F). Error bars denote standard deviation around the mean, significance determined by t-test (n.s. – $p>0.05$, ** - $p<0.01$, *** - $p<0.001$, **** - $p<0.0001$).

**Commented [KB9]:** reword

**Commented [KB10]:** Include description of panel F, with info on number of replicates, and what bars and error bars represent (mean ±SD or SEM)

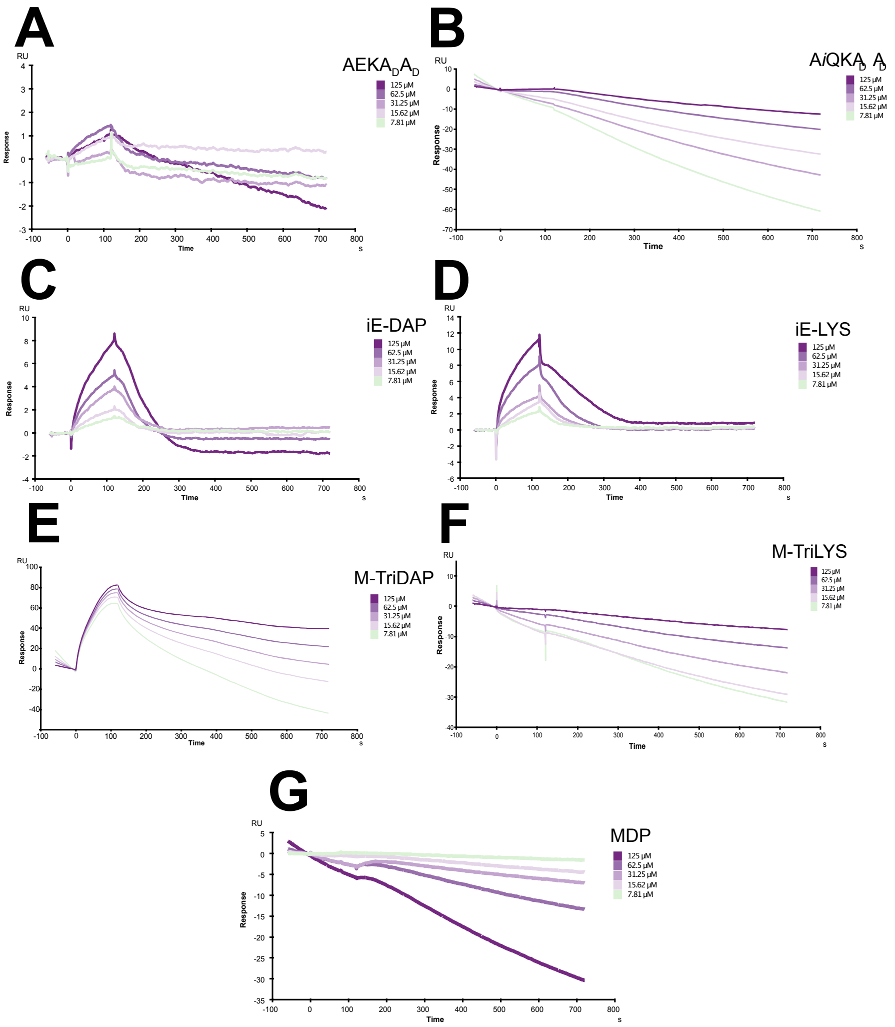


**Supplementary Figure 6 – Ligand binding assessments via SPR to PcsB$_{CHAP}$.** Example sensorgram sets used for kinetic analysis of ligand binding to PcsB$_{CHAP}$ via SPR. Each concentration of analyte was injected onto the sensor chip, presenting the captured PcsB$_{CHAP}$ for 120 s, immediately followed by 600 s injection of running buffer to let ligand and analyte dissociate. The sensorgrams show association and dissociation period of the analysis.

**Commented [KB11]:** Please reword, this is not clear

**Commented [KB12]:** Please revise for readability. Also include information on repeats

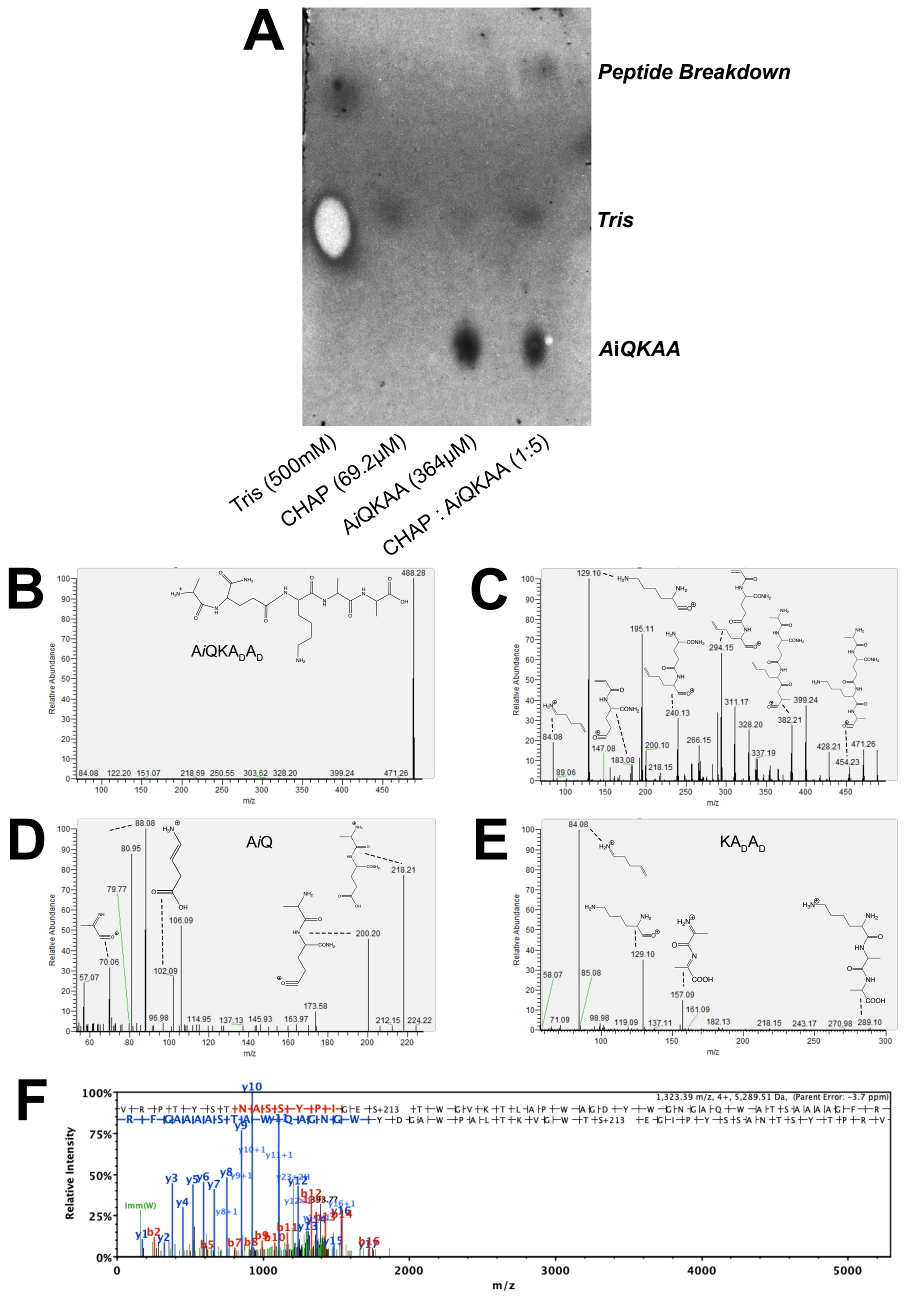


**Supplementary Figure 7 – TLC & Mass Spectrometry evidence for PcsB$_{CHAP}$ Scissile Bond.** TLC of reaction products using water:acetic acid:n-butanol (1:1:3) as the mobile phase (A) showing

Commented [KB13]: define

appearance of polar moiety visualised with ninhydrin. MS1 trace of full synthetic pentapeptide (B) alongside MS2 high energy fragmentation trace (C) confirming species assignment to the full pentapeptide. MS2 traces of isolated product fragments 218m/z (A*i*Q) (D) and 289m/z (KA$_D$A$_D$) (E). Structure of peak shown next to peak. Peak assignment based on predicted m/z readout for PcsB$_{CHAP\ C292S}$ protein (F). A shift of +213 indicates the difference in m/z value for the A*i*Q dipeptide trapped as a covalent intermediate.

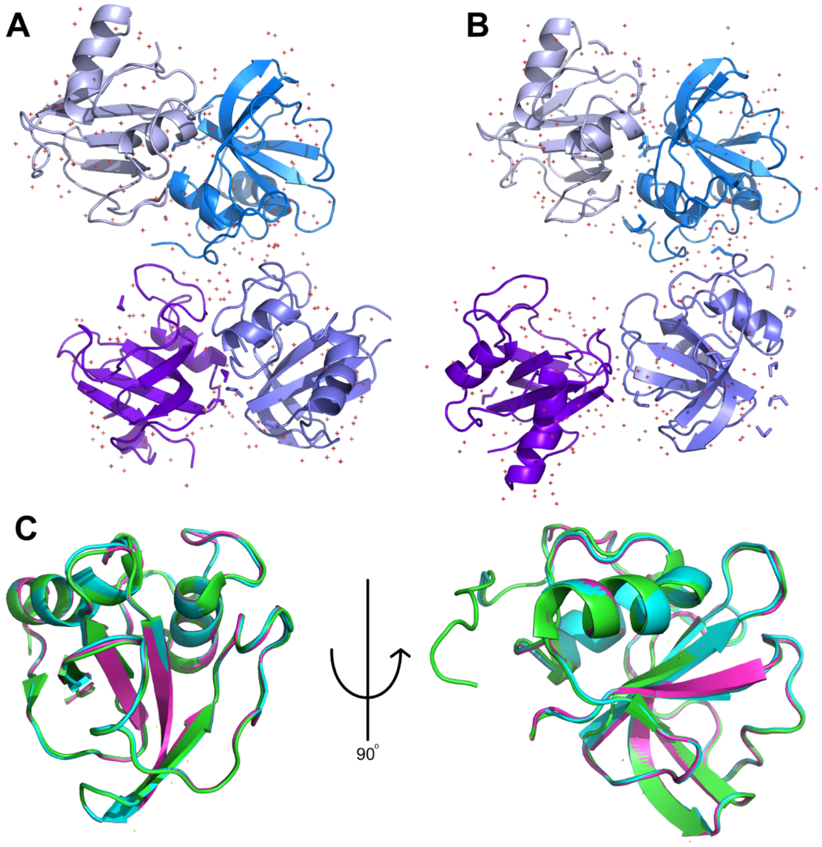


**Supplementary Figure 8 – Crystallographic investigation of PcsB$_{CHAP}$.** X-ray crystal solution structure of PcsB$_{CHAP}$ WT at 1.7 Å (PDB: 9R6W) (A) alongside PcsB$_{CHAP\ C292A}$ at 1.8 Å (PDB: 9R7G) (B). Overlaid structures of CHAP domain from published structure (22) (green, taken from structure of full-length PcsB), WT (cyan) and C292A (pink) (C) demonstrating no obvious changes in structure between

$PcsB_{CHAP}$ and PcsB WT. Red asterisks represent water molecules, whilst short, unconnected chains represent ethylene glycol molecules.

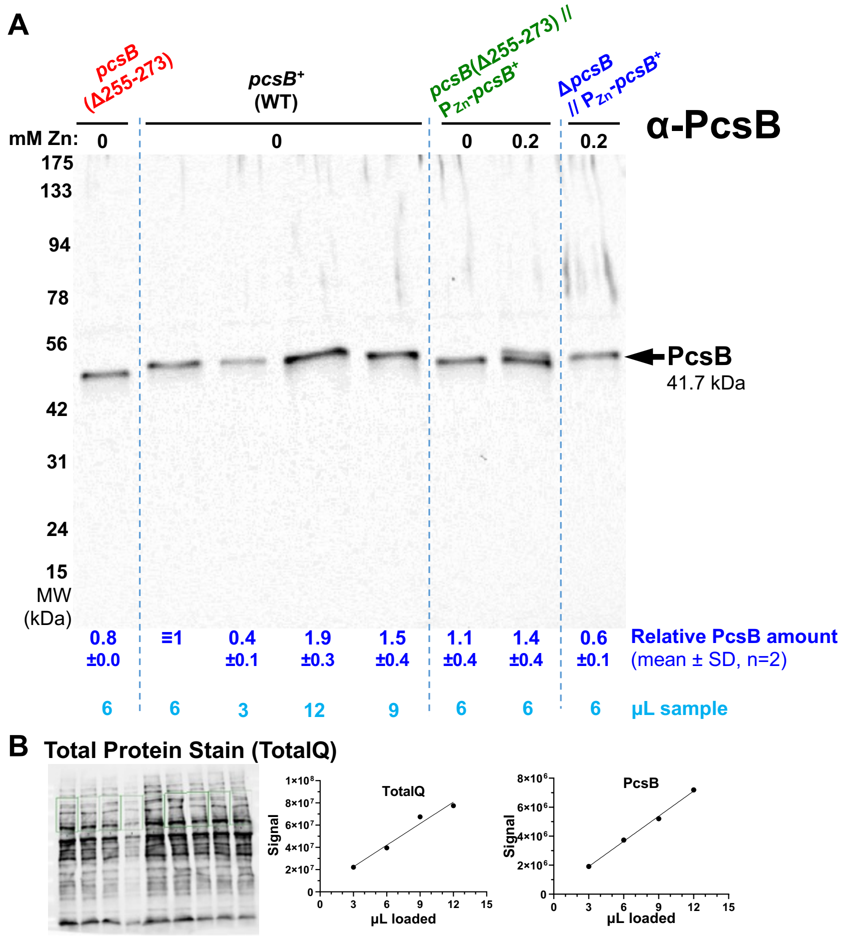


**Supplementary Figure 9—The *pcsB*(Δ255-273) mutant expresses PcsB(Δ255-273) at levels comparable to WT PcsB+ in pneumococcus.** Western blot (A) of pneumococcal cell lysates of *pcsB*(Δ255-273) (IU20481), WT (IU1824), *pcsB*(Δ255-273) // CEP::$P_{Zn}$-*pcsB*+ (IU20370), and Δ*pcsB* // CEP::$P_{Zn}$-*pcsB*+ (IU14806) probed with antibody to native PcsB (see *Methods* for details on antibody generation). (B) The same blot stained with total protein stain, and standard curves used to interpolate values for quantification of protein amounts. Cells were grown in C+Y with Zn inducer as indicated. PcsB amount relative to WT is shown in dark blue (mean ± standard deviation from two biological replicates). Amount of sample loaded in each lane shown in light blue. A representative blot from 2 independent biological replicates is shown.

**Commented [KB14]:** Please replace with the updated version of figure (which I'm sending)

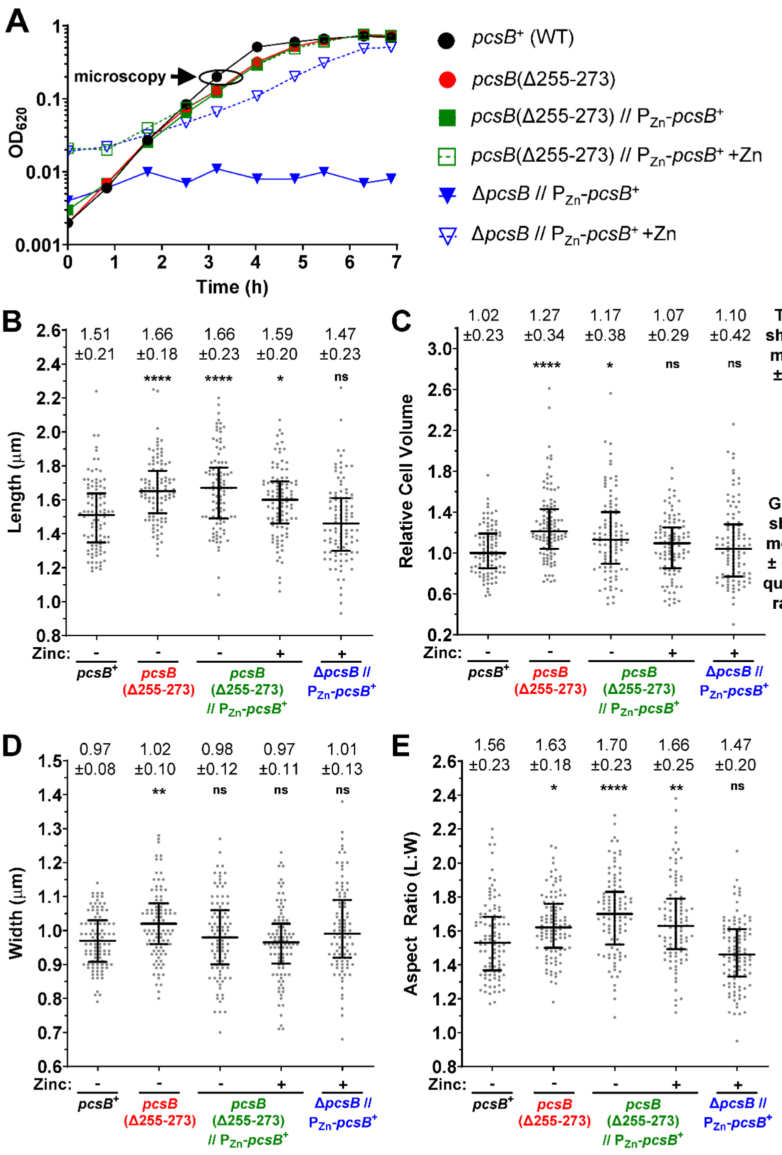


**Supplementary Figure 10 – Removal of the 255-273 helix in PcsB increases cell size but does not affect growth.** Growth curve comparison of WT and *pcsB* mutant pneumococcal strains (A). Pneumococcal cells were grown in C+Y ±0.2 mM Zn inducer and imaged using phase contrast microscopy. From these micrographs, cell length (B), relative volume (C) width (D) and aspect ratio (E) was determined for over 100 cells for each strain from 2 biological replicates. Strains: *pcsB*$^{+}$ (WT, IU1824), *pcsB*(Δ255-273) (IU20481), *pcsB*(Δ255-273) // CEP::P$_{Zn}$-*pcsB*$^{+}$ (IU20370), and Δ*pcsB* // CEP::P$_{Zn}$-*pcsB*$^{+}$ (IU14806). Box plots represent median ± interquartile range, grey dots indicate individual cell measurements. Mean ± SD are indicated along the top. Comparisons between strains were made using a Kruskal-Wallis ANOVA analysis with Dunn's multiple comparisons test. Significance relative to WT is shown. *ns*, not significant; * $P < 0.05$; ** $P < 0.01$; **** $P < 0.0001$.

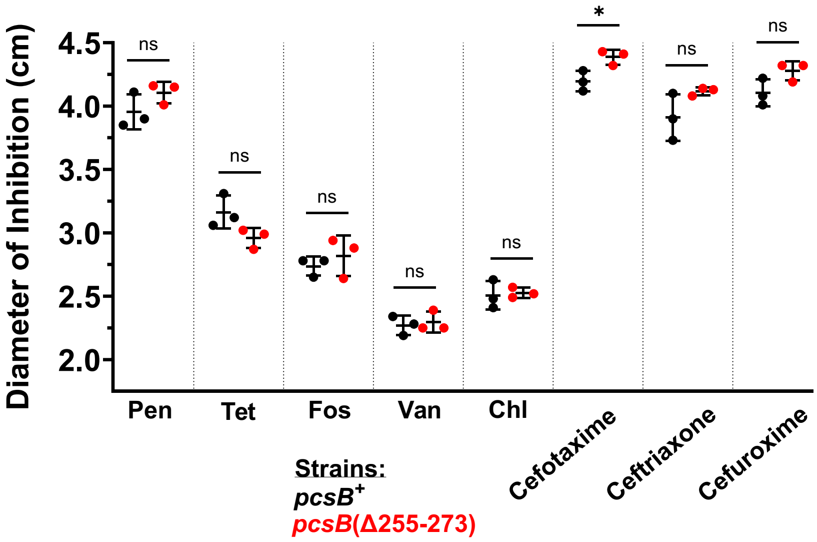


**Supplementary Figure 11 – The pneumococcal *pcsB*(Δ255-273) mutant is more sensitive to cefotaxime, but not other antibiotics tested.** Quantification of the diameter of inhibition for *pcsB*$^{+}$ (WT, IU1824, black) and *pcsB*(Δ255-273) (IU20481, red) pneumococcal strains against penicillin (Pen), cefotaxime, tetracycline (Tet), fosfomycin (Fos), vancomycin (Van), ceftriaxone, chloramphenicol (Chl) and cefuroxime in a disc-diffusion assay. Bars represent means ± standard deviation, with individual replicate values shown as dots. An unpaired *t* test was used to compare between strains, with *p* values shown.
